# Defining phospho-Rab signalling networks downstream of LRRK2 kinase

**DOI:** 10.64898/2026.11.22.767923

**Authors:** Eve Napier, Matthew Jaconelli, Jaijeet Singh, Toan K. Phung, Dario R. Alessi, Amir R. Khan

## Abstract

Activating mutations in leucine-rich repeat kinase 2 (LRRK2) are among the most common genetic risk factors for Parkinson’s disease. LRRK2 phosphorylates a subset of Rab GTPases at a conserved Ser/Thr residue within the switch 2 α-helix, disrupting interactions with canonical regulators and effectors while promoting recruitment of phospho-specific effectors, including RILPL1, RILPL2, JIP3 and JIP4. However, the full spectrum of phosphorylation-dependent Rab interactions remains unknown. Here, we used mass spectrometry to characterise the phospho-Rab interactome. Purified pRab1, pRab3, pRab8, pRab12, pRab35 and pRab43, together with pRab7, a substrate of the related kinase LRRK1, were used as baits in cellular pulldowns. Comparison with non-phosphorylated Rab interactomes identified numerous phosphorylation-dependent binding proteins, alongside known and candidate effectors that preferentially bound non-phosphorylated Rabs, consistent with phosphorylation broadly disrupting Rab complexes. A prominent phospho-selective interactor was TBC1D2 which bound to multiple phosphorylated Rabs. TBC1D2 is a Rab7 GTPase-activating protein (GAP) implicated in LRRK1-dependent regulation of endolysosomal dynamics. AlphaFold modeling, mutagenesis, co-immunoprecipitation and biophysical analyses demonstrated that a conserved di-lysine motif within a central coiled-coil domain of TBC1D2 mediates binding to pRab3A and pRab8A. These findings identify TBC1D2 as a phospho-Rab effector and suggest a molecular link between LRRK2-dependent Rab phosphorylation and Rab GAP function. Our phospho-Rab interaction resource provides a foundation for investigating how LRRK2 regulates membrane trafficking.

## INTRODUCTION

LRRK2 (leucine-rich repeat kinase 2) is a 2,527-amino-acid serine/threonine kinase and the most common genetic contributor to Parkinson’s disease (PD). Mutations in LRRK2 account for approximately 5% of familial and 1% of sporadic PD cases(Healy *et al*, 2008). PD is a progressive neurodegenerative disorder characterised by the selective loss of dopaminergic neurons in the substantia nigra, resulting in debilitating motor and non-motor symptoms(Kalia & Lang, 2015). All established pathogenic LRRK2 mutations, including the common autosomal dominant G2019S variant, enhance kinase activity(Kalogeropulou *et al*, 2022). Moreover, increased LRRK2 kinase activity has also been observed in idiopathic PD(Di Maio *et al*, 2018), raising the possibility that LRRK2-targeted therapies may benefit a broader patient population beyond mutation carriers. Despite intensive investigation and strong genetic validation of LRRK2 as a therapeutic target, the molecular mechanisms by which elevated LRRK2 kinase activity drives cellular dysfunction and ultimately neurodegeneration remain incompletely understood.

LRRK2 phosphorylates a subset of Rab GTPases, including Rab1, Rab3, Rab8, Rab10, Rab12, Rab35 and Rab43, at a conserved serine or threonine residue located within the switch 2 α-helix (e.g. Thr72 in Rab8 and Ser106 in Rab12)(Nirujogi *et al*, 2021; Steger *et al*, 2017; Steger *et al*, 2016). The switch 2 region is a key structural determinant that mediates interactions with regulatory proteins and downstream effectors, suggesting that LRRK2-mediated phosphorylation has the potential to profoundly alter Rab function and signalling.

Rab GTPases constitute the largest family of small GTP-binding proteins and are master regulators of intracellular membrane trafficking in eukaryotic cells(Hutagalung & Novick, 2011). They cycle between an inactive GDP-bound state in the cytosol, and an active GTP-bound state associated with intracellular membranes, where they recruit effector proteins to coordinate vesicle formation, transport, tethering and fusion. This GTP/GDP cycle is tightly controlled by guanine nucleotide exchange factors (GEFs), which promote GTP loading, GTPase-activating proteins (GAPs), which stimulate GTP hydrolysis, and guanine nucleotide dissociation inhibitors (GDIs), which extract Rab proteins from membranes and maintain them in a soluble cytosolic pool(Homma *et al*, 2021; Lamber *et al*, 2019).

Phosphorylation of Rab GTPases by LRRK2 within the switch 2 region has profound functional consequences. Phosphorylated Rab proteins exhibit markedly reduced affinity for guanine nucleotide dissociation inhibitors (GDIs), impairing their extraction from membranes and disrupting normal Rab recycling dynamics. In the case of Rab8, phosphorylation abolishes binding to its cognate guanine nucleotide exchange factor (GEF) Rabin8 and several canonical effector proteins, including OCRL, while preserving interactions with others such as MICAL (Steger et al., 2016; Steger et al., 2017; Waschbüsch et al., 2020). Consequently, phosphorylated Rab proteins become trapped on intracellular membranes, preventing their normal cycling between membrane-associated and cytosolic pools. Consistent with this model, LRRK2-mediated phosphorylation of Rab8 and Rab10 perturbs multiple endolysosomal trafficking pathways, including cargo recycling and membrane transport processes(Rivero-Ríos *et al*, 2020; Rivero-Ríos *et al*, 2019).

Importantly, while phosphorylation disrupts several established Rab interactions, it can also generate novel phospho-dependent binding surfaces that recruit previously unrecognised effector proteins. Recent studies have identified the RILP homology 2 (RH2) domain as a phospho-Rab recognition module that selectively binds phosphorylated Rab8 and Rab10 (Khan et al., 2021; Steger et al., 2017; Waschbüsch et al., 2020). Several cytoskeletal adaptor proteins, including RILPL1, RILPL2, JIP3 and JIP4, contain RH2 domains and are recruited to membranes by phospho-Rab proteins, thereby subverting normal Rab-dependent trafficking pathways (Khan *et al*, 2021; Lara Ordónez *et al*, 2019; Lin *et al*, 2025). These phospho-Rab interactions have important cellular consequences. Binding of RILPL1 to phospho-Rab10 suppresses primary ciliogenesis (Dhekne *et al*, 2018; Lara Ordónez *et al*., 2019; Sobu *et al*, 2021), disrupting a neuroprotective signalling circuit between dopaminergic and cholinergic neurons (Khan *et al*., 2021). Phospho-Rab10–RILPL1 complexes also impair centrosome cohesion during cell division (Fdez *et al*, 2022). JIP3 and JIP4 function as kinesin- and dynein-associated adaptor proteins that regulate intracellular transport. Recruitment of JIP4 by phospho-Rab10 redirects the adaptor to autophagosomes, resulting in aberrant kinesin activation and defects in autophagosome transport and maturation (Boecker *et al*, 2021). In addition, phospho-Rab10-dependent recruitment of JIP4 promotes lysosomal tubulation and vesiculation during lysosomal stress responses (Bonet-Ponce *et al*, 2020). Together, these findings establish that LRRK2-mediated Rab phosphorylation remodels Rab signalling networks by disrupting canonical interactions and promoting the formation of novel phospho-dependent effector complexes. Given that relatively few phospho-Rab effectors have been identified, systematically mapping these interactions could uncover new mechanisms linking pathogenic LRRK2 signalling to altered membrane trafficking, organelle homeostasis and neuronal dysfunction in Parkinson’s disease.

Despite these advances, the phospho-specific interactomes of LRRK2-regulated Rab GTPases remain largely unexplored. Identifying proteins that selectively recognize phosphorylated or non-phosphorylated Rab proteins is therefore critical for understanding the downstream signalling pathways activated by pathogenic LRRK2. To address this question, we developed a toolkit of purified phosphorylated Rab proteins, including pRab1, pRab3, pRab8, pRab12, pRab35 and pRab43, and employed these as affinity reagents in pulldown experiments using A549 cell lysates and mouse brain extracts. pRab7A which is phosphorylated by LRRK1 at the equivalent switch 2 residue Ser72(Hanafusa *et al*, 2019), was also included. Quantitative mass spectrometry was then used to systematically define and compare the interactomes of phosphorylated and non-phosphorylated Rab proteins. This approach identified numerous phospho-Rab-selective binding proteins, including the Rab7A GAP TBC1D2, a protein previously implicated in LRRK1-dependent endosomal regulation(Toyofuku *et al*, 2015). This approach allowed us to explore a potential connection between LRRK2-dependent Rab phosphorylation and the machinery controlling Rab7A activity. More generally, our phospho-Rab interactome platform provides a powerful resource for systematically dissecting how LRRK2-mediated Rab phosphorylation rewires cellular signalling pathways and contributes to Parkinson’s disease pathogenesis.

## RESULTS

### Purification of phosphorylated Rab GTPases

To systematically identify phospho-Rab binding proteins, we first expressed and purified milligram quantities of FLAG-tagged Rab GTPases, missing their C-terminal hypervariable domains (**Fig 1**). LRRK2 substrates, Rab1, Rab3, Rab8, Rab12, Rab35 and Rab43, together with Rab7A, a physiological substrate of LRRK1(Hanafusa *et al*., 2019). To stabilise the GTP-bound conformation, all constructs carried a switch 2 Gln→Leu mutation that is widely used in the Ras superfamily(Scheffzek *et al*, 1997). We omitted the LRRK2 substrate Rab10 from this analysis since it could not be expressed and purified in sufficient quantities for biochemical studies. We also omitted Rab29 which is unusual among Rab GTPases since it predominantly exists in the GDP-state in cells(Gomez *et al*, 2021). We were unable exploit the Gln→Leu variant of Rab29 to stabilize the GTP complex. Previous studies demonstrated that the switch 2 Gln→Leu mutation, which was used to stabilise the GTP-bound state of other Rab proteins, causes Rab29 to misfold (Gomez *et al*., 2021).Consequently, neither Rab10 nor Rab29 were incorporated into the phospho-Rab interactome platform described here.

**Figure 1:**
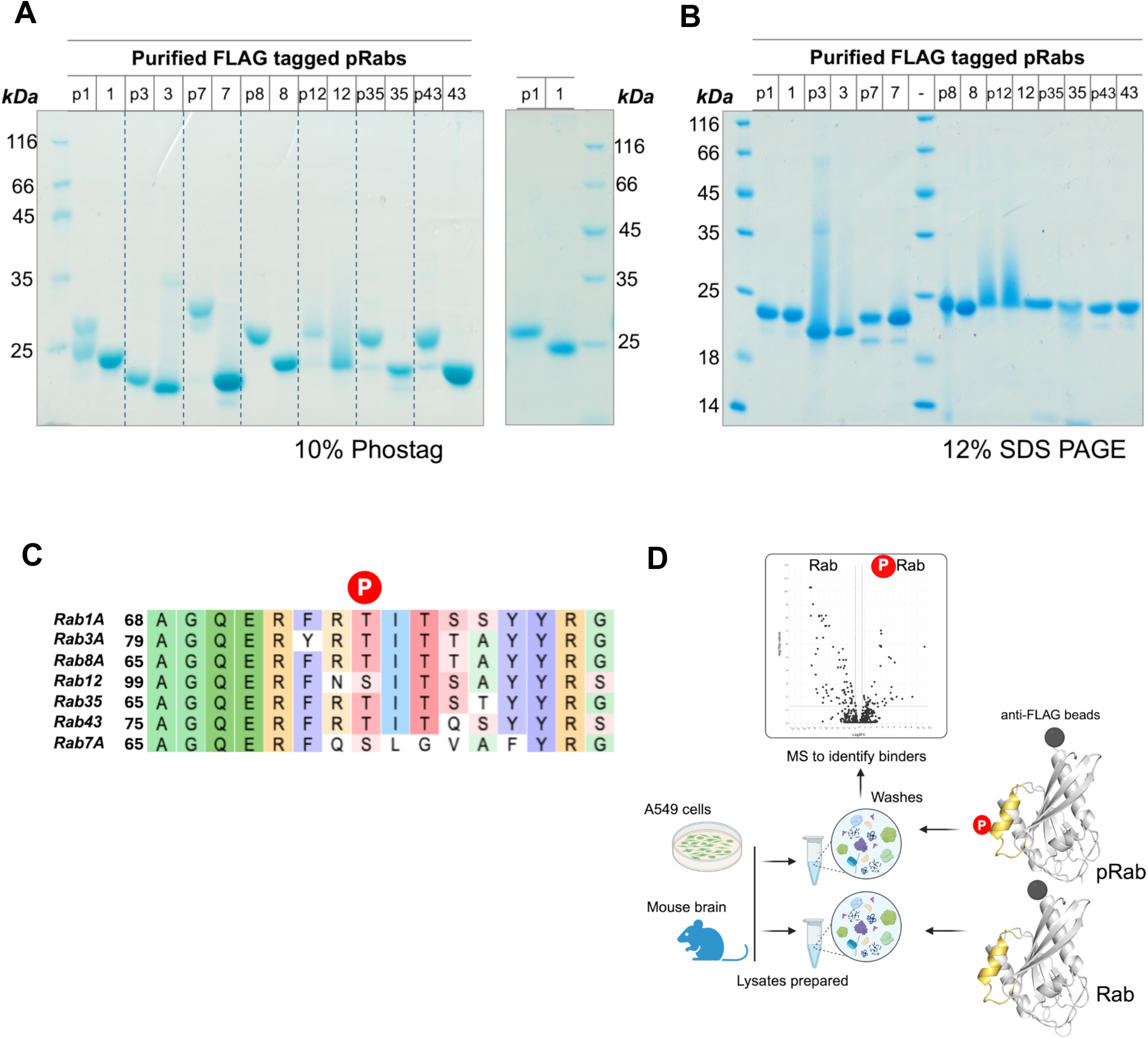
Milligram scale purification of a phospho-Rab toolkit. **(A)** 10% PhosTag gel of the purified proteins. Phosphorylated Rab1A (p1) tends to degrade upon frequent freeze/thaw cycles, therefore a fresh preparation is also shown. **(B)** Conventional SDS-PAGE analyses of the purity of phosphorylated Rab proteins. All Rabs were phosphorylated by MST3, except Rab7 which utilized the enzyme TAK1 kinase. **(C)** Sequence alignment of the switch 2 region of Rabs from our toolkit of phosphorylated proteins. The red P indicates the location of the phosphoserine or phosphothreonine. Alignment was performed using Clustal(Madeira *et al*, 2024), and color annotation used the MAE residue similarity scheme on the web server. **(D)** Work flow to identify phospho-Rab interactors. FLAG-tagged phospho-Rabs were incubated with lysates from either A549 cells or whole mouse brain. Non-phosphorylated FLAG-Rabs were also subjected to the work flow in parallel. Following washes, bound proteins to anti-FLAG beads were analysed by mass spectrometry (example of a volcano plot shown).

Purified Rab proteins were phosphorylated were phosphorylated in vitro at their conserved Ser/Thr residue in switch 2, using a previously established *in vitro* protocol that enabled a crystal structure between pRab8A and its phospho-specific effector RILPL2 (Waschbüsch *et al*, 2020). Minor optimization of the phosphorylation reaction and the purification protocol was performed for each Rab protein (see **Materials and Methods**). We used MST3 (STK24) kinase to phosphorylate the LRRK2 substrate Rabs, as it can be produced in large quantities and modifies all of these Rabs at high stoichiometry on the conserved switch 2 residue targeted by LRRK2(Khan *et al*., 2021; Vieweg *et al*, 2020). The sole exception was Rab7A, which MST3 did not efficiently modify; we therefore phosphorylated Rab7A on Ser72, the site targeted by LRRK1, using TAK1 kinase, which modifies this residue efficiently(Babur *et al*, 2020). High phosphorylation stoichiometry was confirmed by Phos-tag gel electrophoresis, while mutagenesis of the phospho-sites of Rabs provided evidence for correct residue modification by MST3 and TAK1 (**Fig 1 and Suppl Fig S1**). Mutagenesis of FLAG-tagged Rab1A, Rab3A, Rab7A, Rab12, Rab35 and Rab43 at their phospho-sites to a glutamate abolished kinase mediated phosphorylation as evidenced by loss of the singly phosphorylated species (**Suppl Fig S1**). To identify the phospho-site of each phosphorylated Rab, Electron transfer higher collisional dissociation (EThcD) was employed on peptides produced by tryptic digestion of Rab proteins(Penkert *et al*, 2019). This direct strategy confirmed that MST3 phosphorylates the expected switch 2 residue targeted by LRRK2, and in the case of TAK1 kinase, Ser72 of Rab7A (see **Materials and Methods**). Altogether, these efforts established a toolkit of purified FLAG-tagged phospho-Rab proteins comprising pRab1A, pRab3A, pRab7A, pRab8A, pRab12, pRab35 and pRab43. This toolkit formed the basis for the subsequent systematic identification of phosphorylation-dependent Rab-interacting proteins. As a control we employed the non-phosphorylated forms of these proteins.

### Pulldowns of FLAG-Rabs from cell lysates and mass spectrometry

FLAG-tagged Rabs, in both their phosphorylated and non-phosphorylated forms, were used as bait to capture interacting proteins from mouse brain and A549 cell lysates. To lower the levels of endogenous phosphorylated Rabs, and thereby minimise pre-existing phospho-Rab–effector complexes that could compete with the bait, A549 cells were treated with the LRRK2 inhibitor MLi-2 (100nM, overnight) and mice were injected with MLi-2 (30mg/kg, 2h) prior to lysis. Brain tissue and A549 cells were then homogenised in a buffer containing 0.5% detergent together with protein phosphatase inhibitors to inactivate PPM1H and PPM1M in order to prevent dephosphorylation of the pRab proteins. The workflow is summarized schematically in **Fig 1D**. In brief, 400 µg A549 lysate or 750 µg brain homogenate was incubated with 20µg of phosphorylated or non-phosphorylated Rab protein pre-conjugated to 5µL of anti-FLAG magnetic beads. After 1 hour, the beads were isolated, washed twice with high salt lysis buffer (500mM NaCl), followed twice by standard lysis buffer (150mM NaCl). The captured proteins were digested with trypsin and analysed by quantitative high-resolution data-independent acquisition (DIA) liquid chromatography–tandem mass spectrometry (LC-MS/MS). As a control, known interactors RILPL1 and RILPL2 were subjected to the pulldown and visualized by immunoblots to confirm the presence of the interactors enriched within the phosphorylated Rab8 fractions (**Suppl Fig S2**).

Data were analysed using DIA-NN version 1.9.2 and six replicates were performed for each sample. A subset of samples were removed from analysis due to technical concerns arising from sample processing - details of these can be found within the PRIDE submission, where both included and excluded sample data have been uploaded. Briefly, samples that demonstrated significantly reduced TIC, suspected SDS contamination or chromatographic shifts as a result of electrospray instability were removed. To analyse these proteomics data, we used the CURTAIN online tool (https://curtain.proteo.info), which generates, for each Rab protein, a volcano plot of the proteins that bind selectively to its phosphorylated or non-phosphorylated form and allows the underlying primary data to be deconvoluted and visualised. Volcano and violin plots derived from the data are displayed for Rab8A (**Fig 2**), and data for all Rabs are available as CURTAIN links (**Suppl Table 1**). Note that brain Rab3A data were recently reported in another study(Chen *et al*, 2026). CURTAIN derived volcano plots of Rab1A, Rab3A (A549 cells), Rab7A, Rab12, Rab35, and Rab43 are shown as **Suppl Figs S3-S7**.

**Figure 2:**
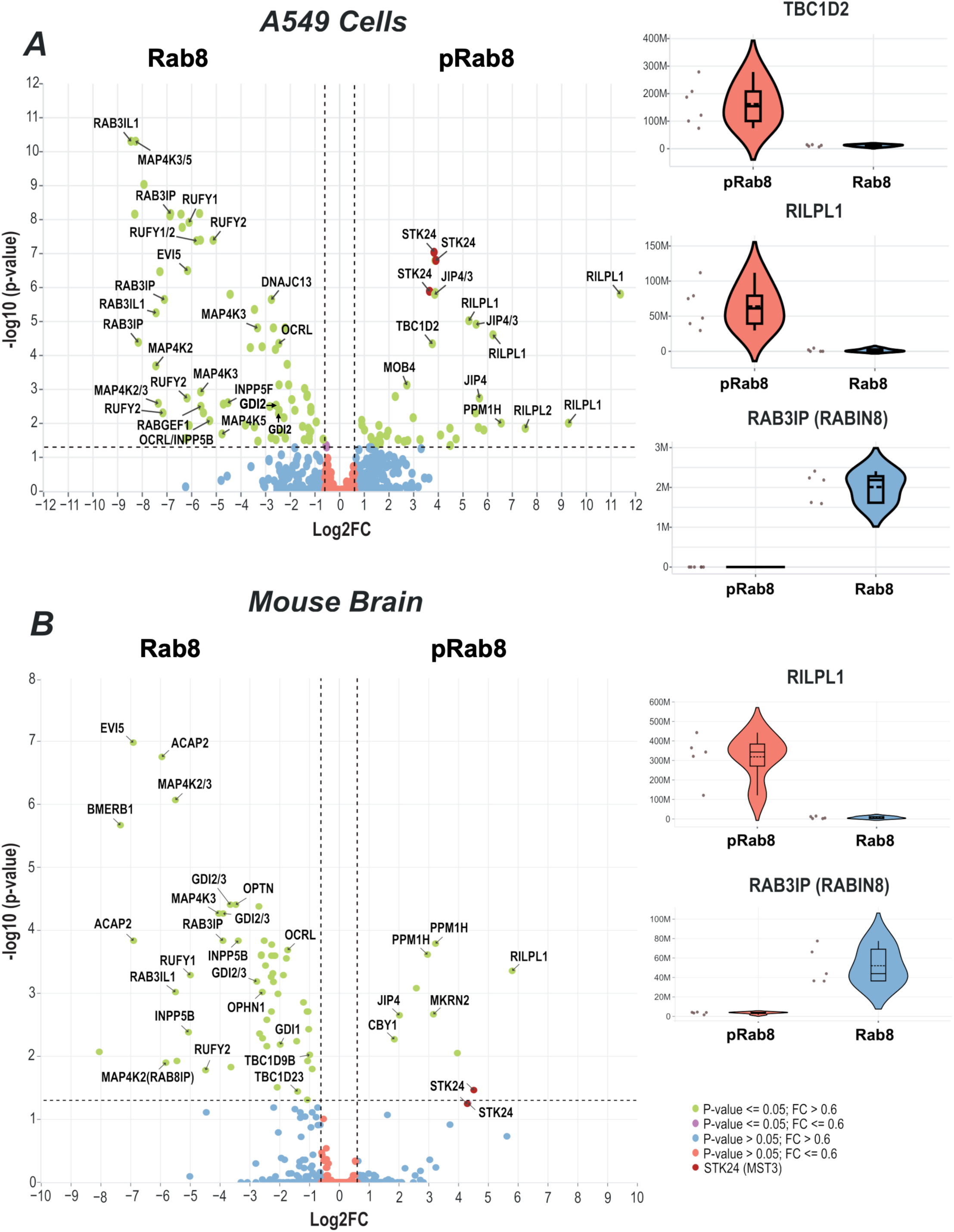
Differential interactome of phosphorylated vs non-phosphorylated Rab8A in A549 cell extract (**A**) and mouse brain homogenates (**B**). Dotted lines indicate Log2FC (vertical) and −log10 p-value (horizontal) cut-offs of 0.6 and <0.05 (1.3; −log10), respectively. Proteins known to associate with Rab8A in a phospho-dependent manner can be observed, such as RILPL1 and RILPL2. Novel interactors significantly enriched in phospho-Rab8A, such as TBC1D2, are additionally annotated. Interactome datasets for each pRab/Rab comparison are freely available using the analysis web tools, Cinder (https://cinder.proteo.info/#/project/39) and Curtain (**Supplementary Table 1**). *Right,* violin plots of known and novel statistically significant (p<0.05) interactors of phosphorylated and non-phosphorylated Rab8A from A549 cells (TBC1D2, RILPL1, Rabin8) and from mouse brain (RILPL1, Rabin8). Intensities are presented as normalised MaxLFQ values from DIA-NN 1.9.2. In panels (A) and (B), STK24 (MST3) is denoted in red, indicating its enrichment as a contaminant for the kinase reaction to generate pRab8A. Some statistically significant hits (green circles, p-value <= 0.05, fold-change (FC) > 0.6) have not been annotated. These proteins have likely been pulled down for technical reasons of the protocol, or they may be nuclear proteins which are unlikely to be genuine interactors of Rab/pRab proteins. However, full annotation of these unlabelled interactors can be found in the CURTAIN links.

**Table 1:** Thermodynamics of pRab8A complexes with TBC1D2 CC domain.

| **K<sub>d</sub> (nM)** | **∆H (kcal/mol)** | **∆S (cal/K•mol)** | **N** |
|---|---|---|---|
| 41 ±30% | -4.9 ±2% | 17 | 1.35 |
| 21 ±50% | -4.3 ±2% | 21 | 1.38 |

For convenience, we additionally built a dedicated tool (https://cinder.proteo.info/#/collate/view/11) that enables a query for any Rab interactor and displays its enrichment across the phosphorylated and non-phosphorylated form of each Rab, in both A549 and brain extracts. As an example, an input of ‘TBC1D2’ in the search query will generate graphs comparing the enrichment of the protein across the full pRab/Rab panel.

Examination of the mass spectrometry data confirmed that, as expected, the known pRab effectors RILPL1 and RILPL2 were enriched with pRab8A in A549 cells (**Fig 2**), a result corroborated by immunoblotting (**Suppl Fig S2**). Other reported pRab effectors were also identified as key hits in the screen, including JIP3 and JIP4 that were enriched with pRab3 and pRab8 in brain and A549 cells (note: identification of JIP3/4 could not be uniquely determined in A549 cells). We also found the phospho-Rab phosphatase PPM1H enriched with pRab8A in both A549 cells and mouse brain. One previously unknown interactor was TBC1D2 (Fig 2A, violin plot), which is investigated in more detail below. A list of selected interactors across the pRab panel, along with their annotations, is provided in **Suppl Table 2**.

Among other known interactors, we found that myosin 5A (Myo5A) binds to both dephosphorylated and phosphorylated Rabs (**Fig 3A**). This may explain why Myo5A is positioned near the centre of volcano plots and why it was not identified as a phospho-selective interactor. Many Rab GTPases including Rab3A, Rab6A, Rab8A, Rab10, Rab11A, Rab14, Rab25, and Rab39B are known to interact with the globular tail domain (GTD) of myosin as a function of gene variants (Myo5A/B/C) that affect the structure of the GTD domain (Lindsay *et al*, 2013; Pylypenko *et al*, 2013; Roland *et al*, 2011; Roland *et al*, 2009). In addition, the adaptor RILPL2 binds to both LRRK2-phosphorylated pRab10 and Myo5A to inhibit ciliogenesis(Dhekne *et al*, 2021). Therefore, the nature and levels of Myo5A/B/C complexes with Rabs be a function of LRRK2 activity, adaptor binding, and Myo5 isoform-specific interactions in various cell types. Rab1A, Rab7A and Rab43 have not previously been identified as Myo5 interactors. However, these findings should be treated with caution since pulldowns from lysates involve the loss of subcellular localizations, and complexes may be indirectly facilitated through common interactors. EHBP1L1 may represent another example of an effector capable of interacting with both forms (**Fig 3B**). EHBP1L1 has previously been reported to interact with Rab proteins including Rab8A/B(Eguchi *et al*, 2018; Nakajo *et al*, 2016), and its enrichment in a phospho-Rab8A pull-down from cells expressing LRRK2 R1441C was reduced following LRRK2 inhibition(Steger *et al*., 2017). In our screens, EHBP1L1 was enriched in pull-downs with pRab3A and pRab12, but no significant differences were found between Rab8A and pRab8A fractions. These findings suggest that EHBP1L1 association with phosphorylated Rab GTPases may depend on the Rab protein and cellular context. Overall these observations raise an important caveat regarding our analyses which were designed to investigate the *enrichment* of the Rab vs. phospho-Rab interactomes. A protein recovered from *both* Rabs and phospho-Rabs will have log2(fold-change) near zero and reside at the centre of the volcano plot. Therefore strong interactors of both Rab and pRab proteins will not be identified with this experimental strategy.

**Figure 3:**
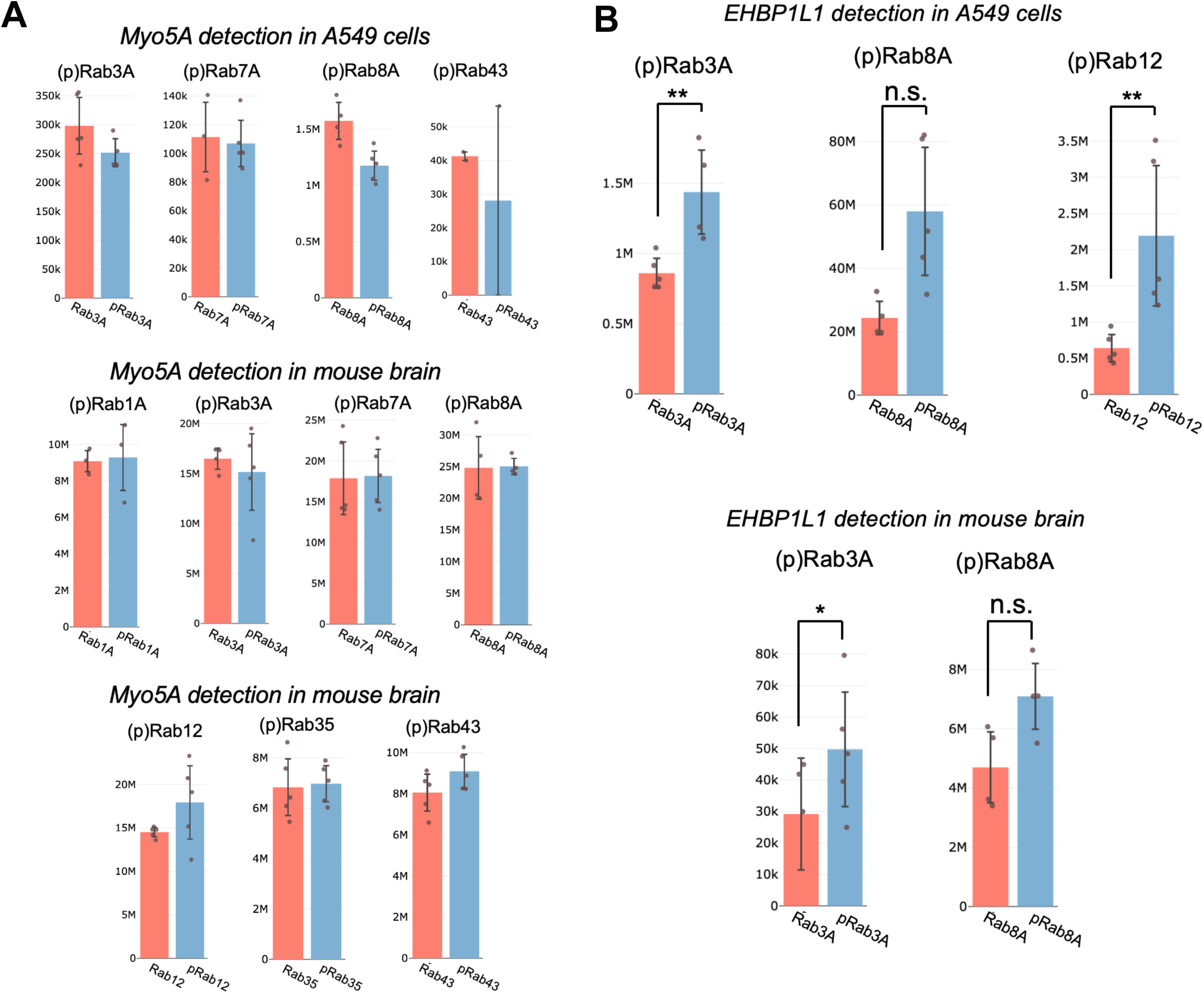
Recovery of known interactors Myo5A and EHBP1L1 from Rab IPs. **(A)** Detection of Myo5A from (p)Rab pulldowns with A549 cell lysates and mouse brain. There was no significant enrichment of Rab or pRab levels in these samples. **(B)** Recovery of EHBP1L1 from (p)Rab pulldowns with A549 and mouse brain lysates. In A549 cells the *p*-values for pRab3A and pRab12 enrichment were 0.006 and 0.007, respectively. There was only marginal enrichment of pRab3A in mouse brain (*p*-value = 0.052), while Rab8A/pRab8A levels had no significant (n.s.) differences (*p*-value > 0.05).

### Rab phosphorylation impairs interaction with many of its effectors

More broadly, our dataset is consistent with the established model that LRRK2-mediated phosphorylation of Rab proteins within the switch 2 region disrupts interactions with canonical Rab regulators and effectors while simultaneously creating binding sites for a smaller number of phospho-selective effectors. Consistent with this, substantially more interactors were identified for non-phosphorylated Rab proteins than for their phosphorylated counterparts (**Suppl Table 1**). Moreover, preferential association with the non-phosphorylated form of a Rab protein provides strong evidence that the interaction is mediated, either directly or indirectly, through the switch 2 region that is modified by LRRK2.

Among the proteins preferentially associated with non-phosphorylated Rab proteins, we identified several established Rab regulators and effectors. For example, Rabin8 and GDI were selectively enriched with non-phosphorylated Rab8A in both A549 cell lysates and mouse brain extracts (**Fig 2**). We previously reported that dephosphorylated Rab3A is preferentially associated with the presynaptic effector proteins RIM1 and RIM2 in mouse brain extracts (Chen *et al*., 2026), and these interactions were markedly reduced following Rab3A phosphorylation. A particularly striking example of phosphorylation-dependent loss of binding emerged from our analysis of Rab43 (**Fig 4**). The TRAnsport Protein Particle (TRAPP) complex exists in two metazoan forms, TRAPPII and TRAPPIII, both of which function as guanine nucleotide exchange factors (GEFs) for Rab43(Harris *et al*, 2021; Jenkins *et al*, 2020). Phosphorylation of Rab43 at Thr82 markedly reduced its interaction with multiple subunits of both TRAPPII and TRAPPIII in mouse brain extracts and A549 cell lysates (**Fig 4**). These findings are consistent with previous observations that the phosphomimetic Rab43[T82D] mutant undergoes substantially slower TRAPPII-mediated nucleotide exchange than wild-type Rab43(Jenkins *et al*., 2020), suggesting that LRRK2-dependent phosphorylation directly impairs activation of Rab43 by its cognate GEF complexes. Together, these data indicate that a major consequence of Rab phosphorylation is the widespread disruption of canonical Rab regulatory and effector interactions, thereby rewiring Rab signalling networks and enabling the recruitment of phospho-selective binding partners. Together, these results show that our workflow for identifying (p)Rab interactors reproduces previously published findings, supporting the validity of the approach.

**Figure 4:**
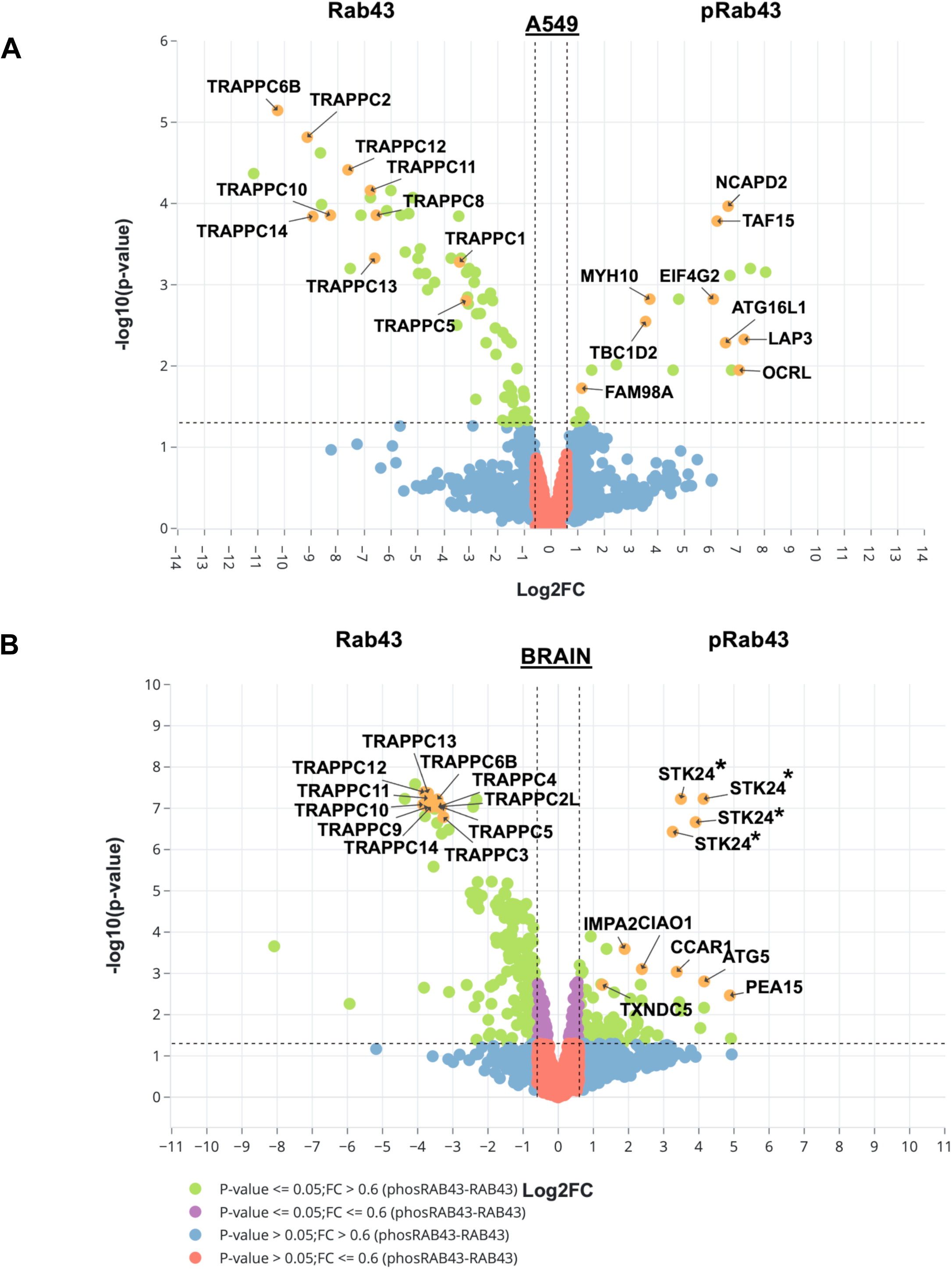
Differential interactomes of phosphorylated vs non-phosphorylated Rab43. Volcano plots show proteins from A549 cell extract **(A)** and mouse brain lysate **(B)**. Dotted lines indicate Log2FC (vertical) and −log10 p-value (horizontal) cut-offs of 0.6 and <0.05 (1.3; −log10), respectively. Proteins are colored according to their respective Log2FC and −log10(p-value). Proteins related to the TRAPPC complex are significantly enriched (p<0.05) within non-phosphorylated Rab43 of both A549 cell extract and mouse brain lysate (colored orange). STK24(MST3) is denoted with asterisk (*) since it is a contaminant from the kinase reaction to generate pRab43.

### TBC1D2 is a novel pRab3A and pRab8A interactor that binds to pRab8A via a central coiled coil domain

Among the proteins most strongly enriched in phosphorylated Rab3A and phosphorylated Rab8A pulldowns was TBC1D2/Armus, a Rab GTPase-activating protein (GAP) of the TBC-domain family. TBC1D2 was enriched in association with all of the LRRK2 phosphorylated Rab proteins (9-fold change for pRab8 in A549 cells; **Fig 5**). In contrast TBC1D2 was not enriched by pRab7A, which is phosphorylated at its switch 2 motif by LRRK1(Hanafusa *et al*., 2019). Intriguingly TBC1D2 has been demonstrated to act as a GAP for the active GTP-bound form of Rab7A (Frasa *et al*, 2010). TBC1D2 is a 928-residue protein that comprises unusual domains for a GAP protein. These include an N-terminal pleckstrin homology (PH) domain (residues 40–150), which has been reported to bind phosphatidylinositol-3-phosphate [PtdIns(3)P] and localise the protein to endosomal membranes; a central coiled-coil domain (residues 300–550), and a C-terminal Tre2/Bub2/Cdc16 (TBC) domain (residues 555–928) that functions as a Rab GTPase-activating protein(Carroll *et al*, 2013; Frasa *et al*., 2010; Law & Rocheleau, 2017; Toyofuku *et al*., 2015)(**Fig 6A**). TBC domains are evolutionarily conserved regulators of small GTPases, and TBC1D2 has previously been shown to function as a GAP for Rab7A and as an effector of Rac1(Carroll *et al*., 2013; Kanno *et al*, 2010). We used Rab8A phosphorylated on Thr72, the physiological LRRK2 phosphorylation site, and predicted its complex with TBC1D2 using AlphaFold modeling(Abramson *et al*, 2024). Confidence models suggested that phosphorylated Rab8A binds to a dimeric form of TBC1D2 through its coiled-coil domain, generating a heterotetrameric complex reminiscent of numerous previously characterised Rab–effector assemblies (Khan & Menetrey, 2013). Details of the modeling protocol and the AlphaFold prediction are provided in **Suppl Fig S8** and in the **Materials and Methods**. The interaction was centred on a conserved basic patch comprising Lys410 and Lys414 located on a predicted α-helix within the coiled-coil domain (**Fig 6B**). In the dimeric model of TBC1D2, these two residues along each α-helix cooperate to engage the phosphate group of a single phospho-Thr72 residue on Rab8A, suggesting a direct phospho-dependent recognition mechanism to from a 2:2 complex. To assess the evolutionary conservation of this predicted binding interface, we performed ConSurf with 405 homologous protein sequences from 394 species obtained from the mammalian dataset in OrthoDB(v13.0). (Yariv *et al*, 2023). Sequences were supplied in FASTA format and aligned using MAFFT, with the evolutionary model setting set to BEST through Contort (https://contort.proteo.nexus/) Both Lys410 and Lys414 were found to be highly conserved (e.g. grade 9 of 9) across mammalian species, consistent with an important functional role in phospho-Rab recognition (**Fig 6C**). Together, these analyses identified a conserved lysine-rich motif within the TBC1D2 coiled-coil domain as a candidate phospho-Rab binding site.

**Figure 5:**
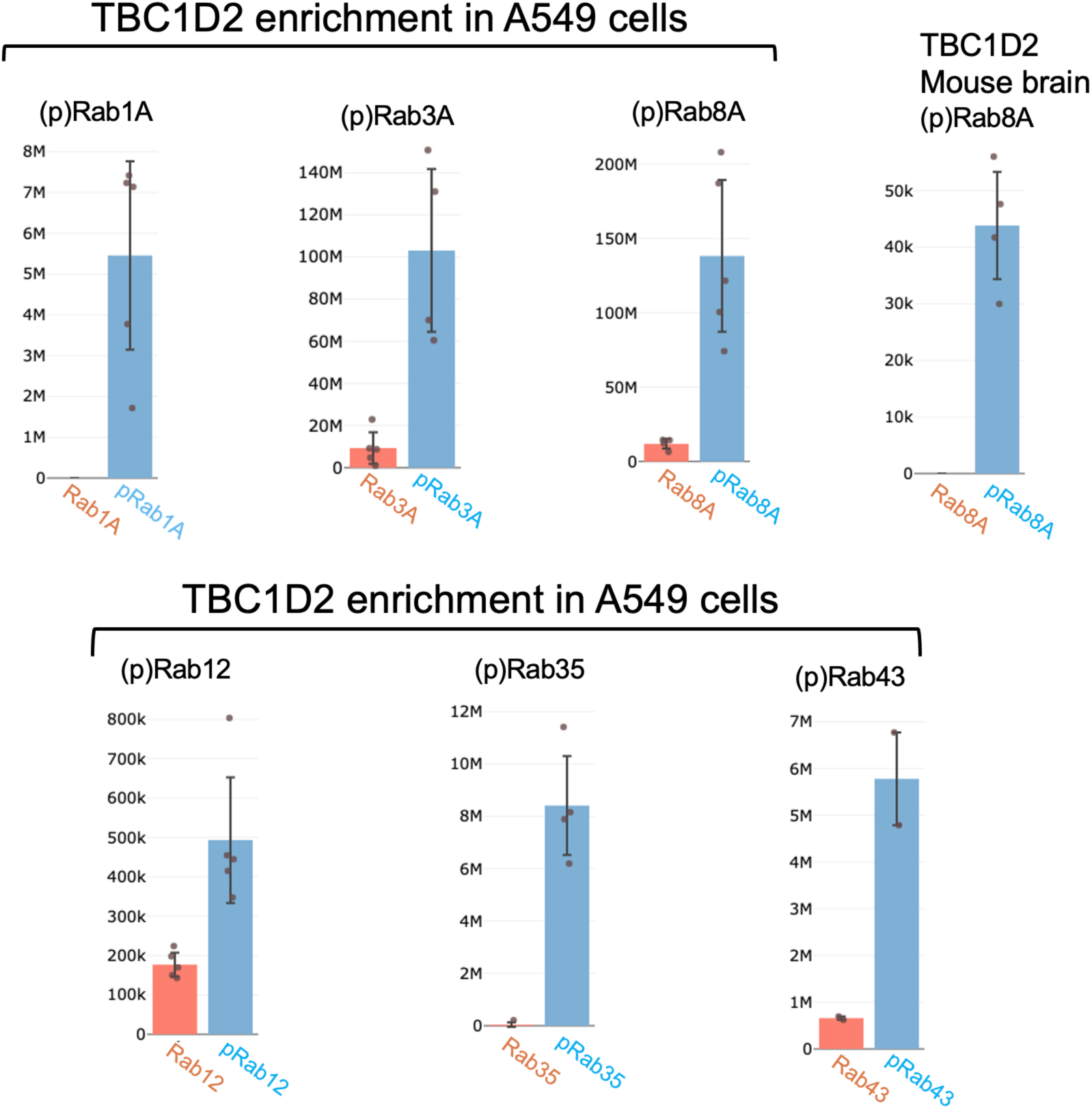
Enrichment of TBC1D2 across the panel of phosphorylated Rab GTPases. TBC1D2 is significantly enriched (*p*-value <0.05) for pRab1A, pRab3A, pRab8A, pRab35 and pRab43, relative to their non-phosphorylated counterparts in A549 cells. There are lower levels of detection for TBC1D2 pulled down with pRab12 in A549 cells, but the enrichment remains significant (*p*-value = 0.001). TBC1D2 pulled down with pRab8a from brain extracts is near the limits of detection.

**Figure 6:**
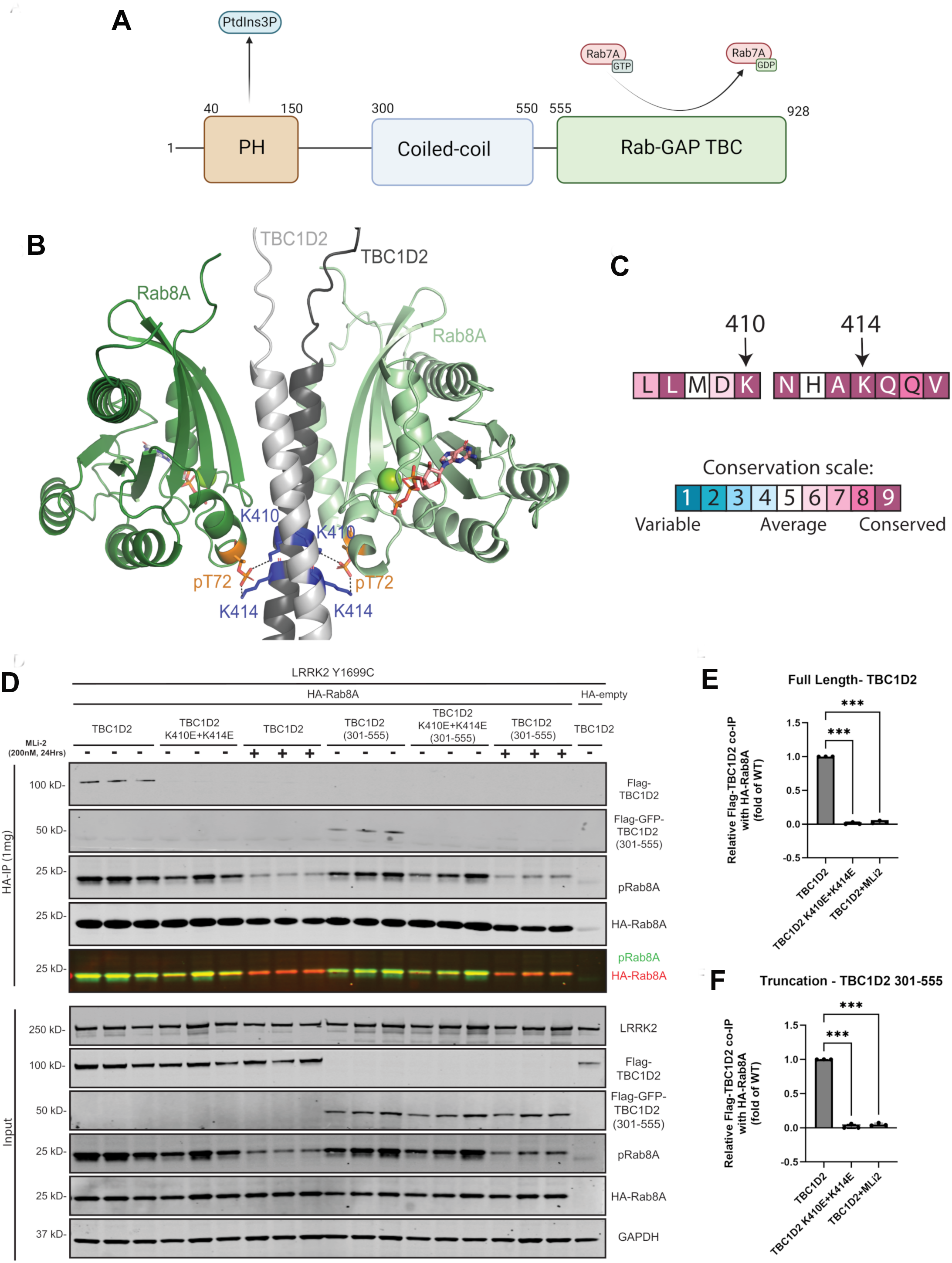
TBC1D2 interacts with pRab8 through a di-lysine motif in its CC domain in HEK293 cells. **(A)** Domain organization of TBC1D2. **(B)** AlphaFold3 model of the interaction between pRab8 and a fragment of the CC domain containing K410 and K414 (blue). TBC1D2 helices are in grey, Rabs in green, phosphothreonine (pT72) is orange, and GTP is shown as a stick model. **(C)** Consurf analysis of the sequence conservation at the Rab-binding region of CC domain. **(D)** Co-immunoprecipitation (co-IP) from HEK293 cells transiently co-expressing hyperactive LRRK2 Y1699C, HA-Rab8A, and either full length FLAG-TBC1D2 or truncation Flag-GFP-TBC1D2 (301-555). Both constructs were tested in their wild-type (WT) and the coiled-coil double mutant (K410E+K414E) forms. Where indicated, cells were treated with the LRRK2 inhibitor MLi-2 (200 nM, 24 h) to block Rab8A phosphorylation. An empty-HA vector served as the negative control for the HA-IP. Immunoblot analysis of anti-HA-Rab8A immunoprecipitation demonstrates that WT TBC1D2 (full length and truncation) specifically co-IPs with phosphorylated Rab8A, whereas this interaction is strongly reduced by either MLi-2 treatment or the K410E+K414E mutation (full length and truncation). **(E & F)** Quantification of full-length TBC1D2 **(E)** and TBC1D2 (301-555) **(F)**. Co-IP TBC1D2 signal was normalised to immunoprecipitated HA-Rab8A and levels of Flag-TBC1D2 in matched input levels. Values are expressed relative to the wild-type control, which was set to 1. Experiments were performed in three independent biological replicates, each comprising three technical replicates (each plotted data point represents the average of the three technical replicates for one biological replicate). Statistical significance was assessed by repeated-measures one-way ANOVA with the Geisser–Greenhouse correction, followed by Dunnett’s multiple-comparisons test (*p < 0.0332, **p < 0.0021, ***p < 0.0002, ****p < 0.0001; ns, not significant).

To experimentally test this model, we established a HEK293 overexpression system in which FLAG-Rab8A, full-length TBC1D2 and the pathogenic hyperactive LRRK2[Y1699C] mutant were co-expressed. Rab8A was subsequently immunoprecipitated and analysed for association with TBC1D2. Under these conditions, robust co-immunoprecipitation of TBC1D2 with Rab8A was observed (**Fig 6D**). Importantly, treatment of cells with the selective LRRK2 inhibitor MLi-2 strongly reduced the interaction, demonstrating that TBC1D2 binding is dependent upon LRRK2-mediated Rab8A phosphorylation (**Fig 6D**). These findings are consistent with the phospho-Rab8A-selective interaction identified in our proteomic screen. To determine whether the predicted basic patch in the coiled-coil domain mediates phospho-Rab8A binding, Lys410 and Lys414 were mutated to glutamate. Strikingly, the K410E/K414E mutant displayed ablated co-immunoprecipitation with Rab8A despite being expressed at levels comparable to wild-type TBC1D2 (**Fig 6D/6E**). Similar results were obtained using a GFP-tagged fragment encompassing the isolated coiled-coil domain of TBC1D2 (residues 301-555). This TBC1D2 variant efficiently associated with Rab8A in cells expressing LRRK2[Y1699C], and the interaction was abolished either by MLi-2 treatment or by introduction of the K410E/K414E mutations (**Fig 6D/6F**). Together, these data support a model in which LRRK2-phosphorylated Rab8A directly engages a conserved lysine-rich motif within the coiled-coil domain of TBC1D2. This previously unrecognised phospho-Rab interaction downstream of LRRK2 may modulate the GAP activity of TBC1D2, its subcellular localization, or both.

### The CC domain is an effector that binds to the GTP conformation of pRab8

We next investigated whether TBC1D2 displays the nucleotide-state selectivity characteristic of bona fide Rab effector proteins. Firstly, we purified pRab8A(GTP), pRab8A(GDP), non-phosphorylated Rab8A(GTP) and a fragment of the CC domain (353-436) fused at the N-terminus to His_6_-NusA (**Fig 7A**). The purity was verified by SDS-PAGE analyses and the hexahistidine-tagged NusA-CC protein was used as bait to pulldown untagged Rab8A variants. *In vitro* pulldown assays revealed that this fragment of the CC domain (353-436) interacts selectively with phosphorylated Rab8A loaded with GTP, whereas no binding was detected to the phosphorylated GDP-bound form (**Fig 7B**). This observation indicates that TBC1D2 preferentially recognises the active, GTP-bound conformation of phosphorylated Rab8A. As a control we performed similar analyses with the RH2 domain of RILPL2, which has previously been shown to possess both GTP and phosphate specificity for pRab8 (**Fig 7C**). Both RILPL2 and the CC domain pulled down pRab8A to similar levels (compare **Fig 7B** and **7C**).

**Figure 7:**
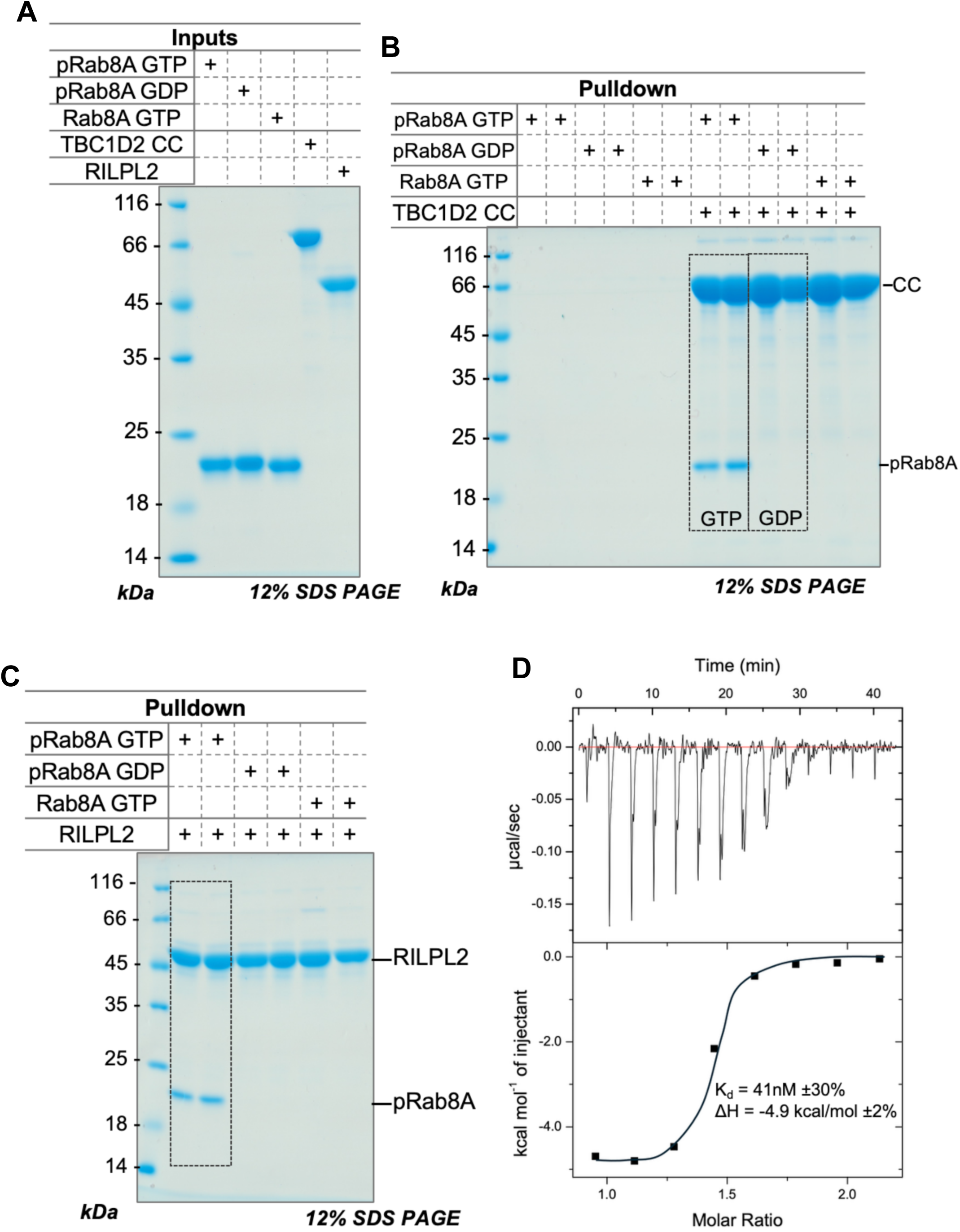
The CC domain of TBC1D2 recognizes the GTP conformation of pRab8A. **(A)** Purified proteins used for the *in vitro* pulldowns, immediately prior to experiments. **(B)** Pulldown of Rab8A variants (prey) with His-tagged CC domain (bait). A fragment of the CC domain (residues 353-436) is a fusion with a His-tag at the N-terminus followed by NusA (His_6_-NusA-CC region). Both the GTP conformation and phosphorylation of Rab8A is required for interactions with this fragment of the CC domain. Rectangular dotted boxes show the difference in pulldown outcomes between GTP and GDP loaded pRab8A. **(C)** Control experiment demonstrating the pulldown of Rab8A with His-tagged RILPL2. The phospho-specific effector RILPL2 pulls down pRab8a to a similar level as the CC region 353-436 of TBC1D2. **(D)** ITC analysis of the interaction between pRab8A (GTP) and a fragment of the CC domain of TBC1D2 (Nus-tagged 353-436). The thermodynamic parameters for this experiment are shown in the figure. Results from two independent experiments are shown in **Table 1**.

To quantify the interaction between the CC domain and pRab8, we performed isothermal titration calorimetry (ITC) using purified proteins. We fused NusA to a more soluble fragment of the CC domain (353-436) to facilitate these experiments. This region of the CC domain nevertheless contained the two Lys residues, K410 and K414, which are predicted to interact with phospho-Thr72 of Rab8A. Two independent titrations were performed yielding similar thermodynamic parameters for the interactions. One representative experiment is shown with a K_d_ = 41nM ±30%, **Δ**H = −4.9 kcal/mol ±2% (**Fig. 7D**). The reported error is calculated from fitting of 8 injections to a sigmoid curve using Origin 7.0 software as described in Protocols.io (https://www.protocols.io/view/isothermal-titration-calorimetry-itc-j8nlkr7x6v5r/v1). The second independent experiment yielded values of K_d_ = 21nM ±50% and **Δ**H = −4.3 kcal/mol ±2%. Raw ITC data have been deposited in the Zenodo server, and the parameters from both experiments are shown in **Table 1**. The higher uncertainty in the K_d_ arises from the steep slope of the sigmoid curve, which lacks sufficient points for more accurate measurements (**Fig 7D**). This curve is typical of high affinity complexes and near the limits of characterisation for the ITC200 calorimeter. In addition, the stoichiometry deviates from 1.0 stoichiometry (inflection point of sigmoidal curve), which would be expected from the AlphaFold prediction of a 2:2 complex (**Fig 6B**). It is important to emphasize that the AlphaFold prediction is marginal in terms of the accuracy of the overall 3-D assembly of TBC1D2 and pRab8A, with a predicted template matching (pTM) score of 0.56 (**Suppl Fig S8**). Scores below 0.50 suggest that predictions of protein complexes are incorrect, while predictions with pTM>0.50 *may be correct* (Abramson *et al*., 2024; Zhang & Skolnick, 2004). Understanding of the stoichiometry and details of the 3-D complex await further structural and biophysical studies – particularly of the full TBC1D2 complex. Although the measured affinity is at the higher end of the range reported for Rab/effector interactions, both the measured K_d_ and enthalpy are consistent with previously characterised Rab/effector complexes (Khan & Menetrey, 2013). Overall, these data establish TBC1D2 as a phospho-Rab effector that selectively recognises the phosphorylated Rab8A and Rab3A through a conserved di-lysine motif within its coiled-coil domain. These findings identify a phospho-Rab-binding motif distinct from the previously characterised RH2 domain of RILP/JIP proteins or the N-terminus of LRRK2.

To further investigate the proposed phospho-Rab8A–TBC1D2 interaction mechanism, we recombinantly expressed and purified multiple TBC1D2 fragments from *E. coli* and assessed their ability to bind phospho-Rab proteins *in vitro* (**Suppl Fig S9**). Pulldown experiments confirmed that the coiled-coil (CC) domain 301-555 of TBC1D2 binds to phosphorylated Rab8A (**Suppl Fig S9A**). In contrast, the core 110-residue PH domain region 43-150 failed to interact. We also expressed a longer version of the PH domain (Nter, **Suppl Fig S9A**) starting from TBC1D2 N-terminus, and which includes a C-terminal region 151-230 that is predicted by AlphasFold3 to interact with the core PH domain. This segment of TBC1D2 1-230 was highly soluble and could be purified to homogeneity, but it also failed to interact with pRab8A (**Suppl Fig S9A**). We also generated single-site and double mutants of the two lysine residues in the CC domain that reversed the positive charges to negatively charged residues (K410E, K414E, and K410E+K414E). In addition, a double mutant K410A+K414A that eliminated both positive charges was generated. None of these mutants of the CC domain had detectable binding to pRab8A (**Suppl Fig S9B**). Thus, both charge reversal and charge neutralisation disrupted the interaction, supporting a critical role for the positively charged lysine residues in phospho-Rab recognition. Altogether, our *in vitro* pulldowns and isothermal titration calorimetry data provide strong support for the interaction model predicted by AlphaFold and are consistent with the proteomic and cellular co-immunoprecipitation analyses.

In the mass spectrometry pull-down experiments using A549 cell lysates, TBC1D2 was enriched with pRab1, pRab3, pRab8, pRab12, pRab35 and pRab43. To examine whether these associations reflect direct binding, we extended the *in vitro* analysis beyond pRab8A to include pRab3A, pRab7A and pRab43. The TBC1D2 CC domain bound pRab3A, but no binding to pRab7A or pRab43 was detected under the conditions tested (**Suppl Fig S9C**). Binding to pRab1, pRab12 and pRab35 was not tested *in vitro*. The discrepancy between pRab43 enrichment in cell lysates and the absence of detectable binding to the isolated CC domain could reflect a weaker interaction or a requirement for other regions of TBC1D2 or additional cellular proteins. Further experiments will be needed to distinguish these possibilities.

## DISCUSSION

Here, we report a system-wide analysis of Rab GTPase interactors regulated by LRRK2-mediated phosphorylation. Beyond identifying phospho-selective binding proteins, our dataset provides a rich resource for defining interactions that are *disrupted* by Rab phosphorylation. Notable examples include the TRAPP trafficking complexes, whose interactions with Rab1A and Rab43 are markedly reduced upon phosphorylation (Rab1 plot, https://cinder.proteo.info/#/analysis-group/180; Rab43 shown in **Fig 4**). There is also a marked reduction in binding of presynaptic RIM1/2 proteins, which recognise Rab3A but not phosphorylated Rab3A(Chen *et al*., 2026). An important future objective will be to generate sufficient quantities of purified, GTP-bound phospho-Rab10 and phospho-Rab29, two physiologically important LRRK2 substrates that could not be analysed here.

Previous studies have mapped Rab interaction networks using GST-Rab pulldowns coupled to mass spectrometry(Gillingham *et al*, 2014; Kanno *et al*., 2010) or proximity-labelling approaches ((Gillingham *et al*, 2019; van Vliet *et al*, 2026). Our strategy uses purified FLAG-tagged phospho-Rab proteins to examine phosphorylation-dependent interactions directly. Recovery of established phospho-Rab interactors, including RILPL1, RILPL2, JIP3 and JIP4, supports the validity of this approach. However, cell lysis can bring together proteins that would not normally encounter one another in intact cells. Interactor recovery also depends on wash stringency, as illustrated by the differences between high- and low-stringency conditions (**Suppl Fig S2**). The resulting datasets therefore capture interactions recoverable under the selected experimental conditions rather than a complete or definitive interactome. Candidate interactors require further cellular and biochemical validation to establish their physiological relevance.

Our study has several limitations. First, the Rab proteins used as baits lacked their C-terminal hypervariable regions and were therefore neither prenylated nor membrane-anchored. Interactions that depend on these regions, their targeting determinants or the membrane environment may consequently be under-represented. Second, all baits carried a switch 2 Gln→Leu substitution to stabilize the GTP-bound state, so interactions requiring nucleotide cycling or the GDP-bound conformation would not be captured. Third, our approach measures differential enrichment rather than absolute binding. Effectors that bind similarly to phosphorylated and non-phosphorylated Rabs may therefore lie near zero on the fold-change axis of the volcano plots and escape identification as phospho-selective interactors. Myosin Va illustrates this limitation, as it binds to both dephosphorylated and phosphorylated Rab8A and Rab10 (Dhekne et al., 2021). EHBP1L1 may provide a further example of an effector whose association is not uniformly promoted by Rab phosphorylation. It is an established Rab effector(Eguchi *et al*., 2018; Liu *et al*, 2020; Nakajo *et al*., 2016) but it was also found to be enriched with immunoprecipitated LRRK2 phosphorylated Rab8A in a previous study(Steger *et al*., 2017). Previous studies have implicated it in LRRK2-dependent lysosomal responses, whereas phosphorylation of Rab10 at Thr73 was reported to reduce its association with EHBP1L1(Eguchi *et al*., 2018; Liu *et al*., 2020). Together with the differing enrichment patterns observed in our screens (**Fig 3**), these findings suggest that the effect of phosphorylation on EHBP1L1 association may depend on the Rab protein and experimental context. A lack of phospho-selective enrichment should therefore not be interpreted as an absence of binding to phosphorylated Rabs. Finally, our datasets derive from a single human cell line and mouse brain tissue. Additional phosphorylation-dependent interactions are likely to occur in other cellular contexts, including defined neuronal and immune-cell populations.

One of the most intriguing findings of this study is the identification of TBC1D2 as a phospho-Rab effector. Using cellular co-immunoprecipitation, *in vitro* pulldown assays, mutagenesis and isothermal titration calorimetry, we show that a conserved di-lysine motif within the coiled-coil domain of TBC1D2 mediates high-affinity binding to phosphorylated Rab8A and Rab3A. This establishes the TBC1D2 coiled-coil domain as a phospho-Rab(GTP) recognition module - only the second such motif to be characterised, alongside the RH2 domains of the RILP-like and JIP protein families.

TBC1D2 has an unusual domain architecture: an N-terminal PH domain that binds PtdIns(3)P, a central coiled-coil domain, and a C-terminal TBC GAP domain active toward Rab7A(Carroll *et al*., 2013; Frasa *et al*., 2010). Notably, Rac1 has been reported to bind residues 295–433 of the coiled-coil region(Frasa *et al*., 2010), that encompasses the K410/K414 pRab binding CC domain. This raises the possibility that phospho-Rab8A and Rac1 compete for TBC1D2, such that pathogenic LRRK2 activity could shift TBC1D2 away from Rac1-dependent functions in cytoskeletal remodelling and cell–cell adhesion(Mori *et al*, 2021).

Our identification of the phospho-Rab8A-binding site raises experimentally testable hypotheses that could be investigated using purified proteins, in vitro GAP assays, and cellular studies. Phospho-Rab8A binding to the coiled-coil domain may modulate the inhibitory interaction between the PH and TBC domains, potentially influencing TBC1D2 GAP activity. TBC1D2 is thought to adopt an autoinhibited conformation maintained by intramolecular interactions between its PH and catalytic TBC domains(Law & Rocheleau, 2017; Toyofuku *et al*., 2015), with Vps34-generated PtdIns(3)P and LRRK1-dependent signalling implicated in its recruitment and activation. Our findings raise two testable possibilities: pRab8A binding could alter the subcellular localisation and availability of TBC1D2 (**Fig 8**). In this model, binding to phospho-Rab8A could redistribute TBC1D2 away from Rab7-positive endosomal and lysosomal membranes, potentially reducing its access to Rab7A and altering Rab7A-dependent trafficking. An alternative model is that engagement of the TBC1D2 coiled-coil domain with pRab8A could alter PH–TBC domain autoinhibition. Relief of this autoinhibitory interaction would be predicted to increase Rab7A GAP activity, thereby inactivating Rab7A. These mechanisms remain hypothetical and will require direct measurements of TBC1D2 localisation, GAP activity and Rab7A-GTP levels. Such studies will determine whether phospho-Rab binding alters TBC1D2-dependent Rab7A regulation and contributes to the endolysosomal defects associated with pathogenic LRRK2 activation. These future studies could potentially establish a point of convergence between LRRK1 and LRRK2 signalling (**Fig 8**).

**Figure 8:**
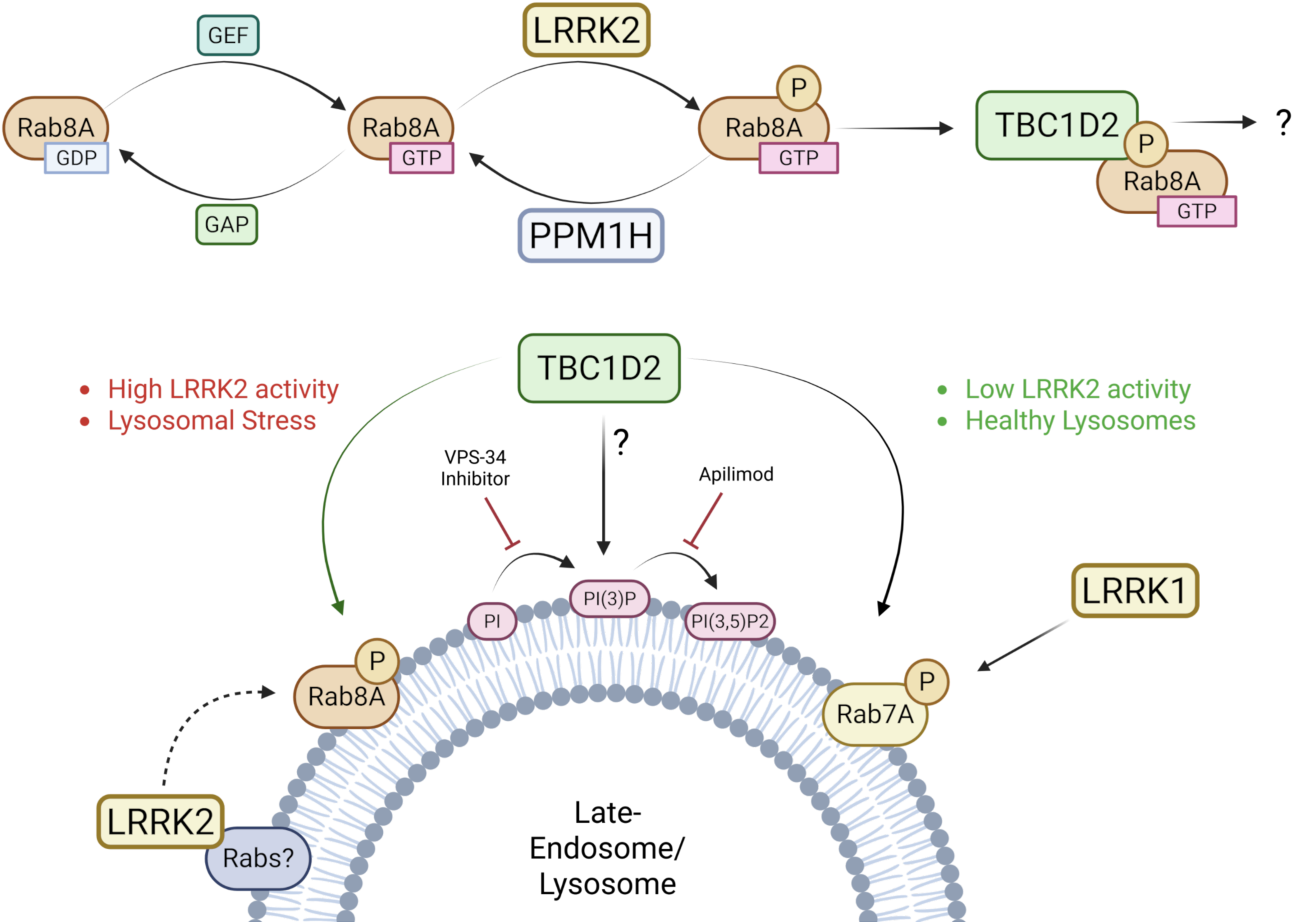
Proposed model linking LRRK2 signalling, TBC1D2 and Rab7A regulation. LRRK2-mediated phosphorylation and PPM1H-mediated dephosphorylation regulate Rab8A phosphorylation. In response to lysosomal stress, increased LRRK2-dependent Rab8A phosphorylation may promote the formation of pRab8A–TBC1D2 complexes. We propose that this interaction could redistribute TBC1D2 away from Rab7A-positive membranes, reducing its access to Rab7A and potentially altering Rab7A-dependent endosomal and lysosomal trafficking. Alternatively, pRab8A binding could modulate TBC1D2 GAP activity by altering PH–TBC domain autoinhibition. LRRK1 has previously been implicated in promoting TBC1D2-dependent Rab7A inactivation and also phosphorylates Rab7A; how these activities relate to the proposed LRRK2–pRab8A–TBC1D2 pathway remains unresolved. Solid arrows indicate supported relationships, including regulation of Rab8A phosphorylation by LRRK2 and PPM1H and the pRab8A–TBC1D2 interaction identified here. Dashed arrows indicate hypothetical steps. Redistribution of TBC1D2, changes in Rab7A-GTP levels and modulation of TBC1D2 GAP activity have not been experimentally established in this study.

Overall, our study establishes a versatile phospho-Rab interactome platform for dissecting signalling networks downstream of LRRK2 and other Rab kinases. The identification of TBC1D2 as a phospho-Rab effector demonstrates the utility of this approach and provides a starting point for investigating how Rab phosphorylation influences Rab GAP function. Extending this analysis to additional Rabs and cellular contexts may reveal further phosphorylation-selective interactions and clarify how pathogenic LRRK2 activity disrupts membrane trafficking in PD.

## Materials and Methods

### Key Resource Table is provided

#### Reagents

MLi-2, a potent and selective LRRK2 inhibitor, was synthesised by and obtained from MRC Protein Phosphorylation and Ubiquitylation Unit (MRC PPU) Reagents and Services, University of Dundee, and is available to request via the MRC PPU Reagents and Services website (https://mrcppureagents.dundee.ac.uk). MLi-2 was dissolved in DMSO to a 10 mM stock and stored at −20 °C. An equivalent volume of DMSO was used as vehicle control in all experiments.

#### Plasmids

All plasmids used in this study encode human proteins and were generated by MRC PPU Reagents and Services, University of Dundee; they are available to request via the MRC PPU Reagents and Services website (https://mrcppureagents.dundee.ac.uk). Full sequences and maps are available under the DU numbers listed below.

#### List of plasmids

| DU number | Construct |
|---|---|
| DU80271 | pCMV5D FLAG–TBC1D2 (full length) |
| DU80507 | pCMV5D FLAG–TBC1D2[K410E/K414E] |
| DU80501 | pCMV5D FLAG–GFP–TBC1D2[301–555] |
| DU80521 | pCMV5D FLAG–GFP–TBC1D2[301–555] K410E/K414E |
| DU35414 | pCMV5 HA–Rab8A |
| DU52369 | pcDNA5D FRT/TO LRRK2[Y1699C] |
| DU49302 | pCMV5 HA (empty vector control) |

#### List of primary antibodies

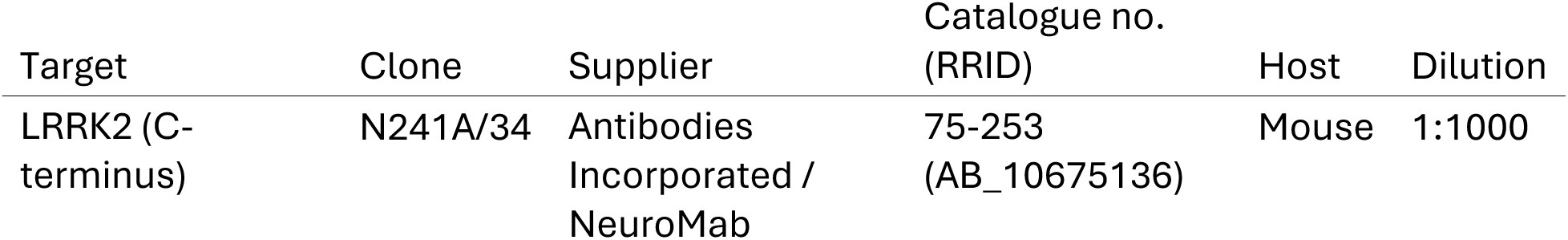
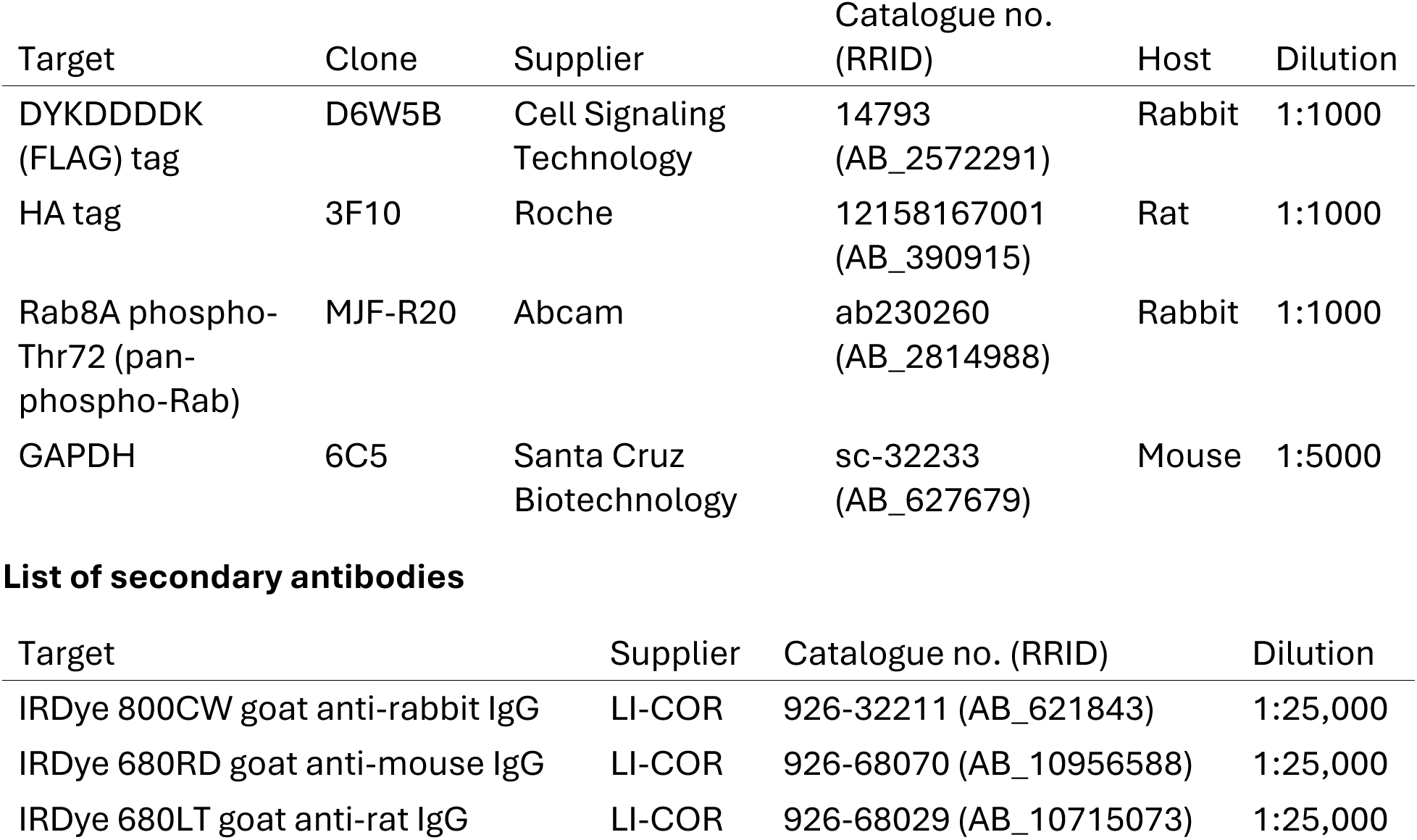

| Target | Clone | Supplier | Catalogue no. (RRID) | Host | Dilution |
|---|---|---|---|---|---|
| LRRK2 (C-terminus) | N241A/34 | Antibodies Incorporated / NeuroMab | 75-253 (AB_10675136) | Mouse | 1:1000 |
| Target | Clone | Supplier | Catalogue no. (RRID) | Host | Dilution |
| DYKDDDDK (FLAG) tag | D6W5B | Cell Signaling Technology | 14793 (AB_2572291) | Rabbit | 1:1000 |
| HA tag | 3F10 | Roche | 12158167001 (AB_390915) | Rat | 1:1000 |
| Rab8A phospho-Thr72 (pan-phospho-Rab) | MJF-R20 | Abcam | ab230260 (AB_2814988) | Rabbit | 1:1000 |
| GAPDH | 6C5 | Santa Cruz Biotechnology | sc-32233 (AB_627679) | Mouse | 1:5000 |
| **List of secondary antibodies** |  |  |  |  |  |
| Target |  | Supplier | Catalogue no. (RRID) |  | Dilution |
| IRDye 800CW goat anti-rabbit IgG |  | LI-COR | 926-32211 (AB_621843) |  | 1:25,000 |
| IRDye 680RD goat anti-mouse IgG |  | LI-COR | 926-68070 (AB_10956588) |  | 1:25,000 |
| IRDye 680LT goat anti-rat IgG |  | LI-COR | 926-68029 (AB_10715073) |  | 1:25,000 |

### Cell culture, transfection and lysis

HEK293 cells (ATCC, CRL-1573; RRID:CVCL_0045) were cultured in Dulbecco’s Modified Eagle Medium (DMEM; Gibco) supplemented with 10% (v/v) foetal bovine serum (Gibco), 2 mM L-glutamine (Sigma-Aldrich), 100 U/mL penicillin and 100 µg/mL streptomycin (Sigma-Aldrich), and maintained at 37 °C in a humidified incubator with 5% CO₂. Cells were confirmed free of mycoplasma contamination by routine testing.

For co-immunoprecipitation experiments, cells were seeded in 10 cm dishes and transfected at 70–80% confluency using polyethyleneimine (PEI; Polysciences). Per dish, 7 µg of pcDNA5D FRT/TO LRRK2[Y1699C], 2 µg of the indicated FLAG–TBC1D2 construct (full-length wild type, full-length K410E/K414E, Flag-GFP–TBC1D2[301–555] or Flag-GFP–TBC1D2[301–555] K410E/K414E) and 1 µg of either HA–Rab8A or the corresponding HA empty vector were diluted in 500 µL Opti-MEM reduced-serum medium (Gibco) and combined with 40 µg PEI (4:1 PEI:DNA, w/w). The complexes were allowed to form for 30 min at room temperature before being added dropwise to the culture medium. Cells were lysed 24 h after transfection. Where indicated, cells were treated with 200 nM MLi-2 or an equivalent volume of DMSO for 24 h prior to lysis.

Cells were washed in ice-cold PBS and lysed in ice-cold buffer containing 50 mM Tris-HCl pH 7.5, 150 mM NaCl, 10mM β-glycerophosphate, 10 mM sodium pyrophosphate, 10% (v/v) glycerol and 1% (v/v) Triton X-100, supplemented immediately before use with one tablet each of PhosSTOP phosphatase inhibitor cocktail (Roche, 04906837001) and cOmplete Mini EDTA-free protease inhibitor cocktail (Roche, 11836170001) per 10 mL of buffer. Lysates were clarified by centrifugation at 17,000 × g for 10 min at 4 °C, and protein concentration in the supernatant was determined by Bradford assay (Bio-Rad Protein Assay Dye Reagent Concentrate, 5000006).

### Immunoprecipitation

Clarified lysate (1 mg of total protein) was incubated with 40 µL of Pierce anti-HA magnetic bead slurry (Thermo Fisher Scientific, 88837) that had been pre-washed in 1× PBS. Samples were incubated for 1 h at 4 °C with gentle rotation at 20 rpm. Beads were collected on a DynaMag-2 magnet (Invitrogen, 10723874) and washed three times with 1 mL of TBS-T (20 mM Tris-HCl pH 7.5, 150 mM NaCl, 0.1% (v/v) Tween-20). Bound proteins were eluted by heating the beads in 30 µL of 2× NuPAGE LDS sample buffer supplemented with 1% (v/v) β-mercaptoethanol at 70 °C for 10 min. Eluates were separated from the beads on the magnet and retained for immunoblotting.

### Quantitative immunoblotting

Immunoblotting was performed essentially as described (protocols.io; https://dx.doi.org/10.17504/protocols.io.ewov14znkvr2/v2). Briefly, 20–25 µg of total cell lysate or 30% of each immunoprecipitated fraction was resolved on precast NuPAGE 4–12% Bis-Tris midi gels (Thermo Fisher Scientific, WG1402BOX) at 120 V in NuPAGE MOPS-SDS running buffer (Thermo Fisher Scientific, NP0001-02). Proteins were transferred to nitrocellulose membranes (Amersham Protran Supported, 0.45 µm; Cytiva) for 90 min at 90 V in transfer buffer (48 mM Tris, 39 mM glycine, 20% (v/v) methanol). Membranes were blocked in 5% (w/v) non-fat dried milk in TBS-T for 1 h at room temperature, then incubated overnight at 4 °C with primary antibodies diluted in 5% (w/v) BSA in TBS-T. Membranes were washed three times for 10 min in TBS-T, incubated for 1 h at room temperature with the appropriate IRDye-conjugated secondary antibody diluted in 5% (w/v) non-fat dried milk in TBS-T, and washed a further three times for 10 min in TBS-T. Fluorescent signals were acquired on a LI-COR Odyssey CLx imaging system and band intensities quantified using Image Studio Lite (version 5.2.5; RRID:SCR_013715).

### Quantification and statistical analysis

All experiments were performed in three independent biological replicates. For co-immunoprecipitation experiments, the amount of co-precipitated protein was normalised to the corresponding immunoprecipitated HA–Rab8A signal and to the level of the FLAG-tagged protein in the matched input, and values were expressed relative to the wild-type control, which was set to 1. Data are presented as mean ± SD. Statistical significance was assessed by repeated-measures one-way ANOVA with the Geisser–Greenhouse correction, followed by Dunnett’s multiple-comparisons test in GraphPad Prism 10.5.0 (RRID:SCR_002798); P < 0.05 was considered significant.

### Expression and purification of Rabs

dx.doi.org/10.17504/protocols.io.m7vznmo54vx1/v1

Rab GTPases were expressed and purified as described in Protocols.io. Briefly, the cDNA for all Rab constructs (FLAG Rab1A 1-176 Q70L, FLAG Rab3A 18-190 Q81L, FLAG Rab8A 1-181 Q67L, FLAG Rab12 36-208 Q101L, FLAG Rab35 1-180 Q67L, FLAG Rab43 9-190 Q77L, FLAG Rab7A 1-176 Q67L,) and MST3 kinase codon optimised for *E.coli* expression, were ordered from Genscript and cloned into pET28a at the NdeI/BamHI sites, transformed into E. coli BL21 DE3 (ThermoFisher Scientific). All Rabs were grown in 2xYT, supplemented with 30µg/ml kanamycin at 37℃. At OD600 ∼ 0.6 was reached, cells were induced with 0.5mM IPTG overnight at 18℃. MST3 was grown in the same conditions in LB broth, and induced with 0.1mM IPTG. The cells were then also incubated overnight at 18℃. Cells were harvested by centrifugation (5000 x g, 10 minutes, 4℃) and lysed by sonication in extraction buffer (20 mM Tris-HCl pH 8, 300 mM NaCl, 20 mM imidazole, 10 mM β-mercaptoethanol, 5 mM MgCl2). Lysates were clarified by centrifugation (20,000 x g, 40 minutes, 4℃) and applied to a Ni2+-agarose resin gravity flow column.

The resin was washed with extraction buffer and wash buffer (extraction buffer supplemented with 40mM Imidazole), and eluted in extraction buffer containing 200mM Imidazole. The His-tag was removed from all FLAG-Rabs by overnight dialysis at 4℃ in gel filtration buffer (20mM Tris-HCl, pH 8.0, 100 mM NaCl, 1 mM DTT, 5 mM MgCl2) with thrombin protease, followed by a second Ni2+-column and concentration using an Amicon® Ultra centrifugal filter. A portion of each Rab (4-5mg) was further purified by gel filtration using a Superdex 75 column (10/300 GL) equilibrated in gel filtration buffer and used as non-phosphorylated controls.

### Phosphorylation of Rabs

https://dx.doi.org/10.17504/protocols.io.m7vznmo54vx1/v1

Rab GTPases were phosphorylated as described in [<u>protocols.io</u>]. It has previously been shown that MST3 kinase can efficiently phosphorylate Rab8A at T72 in vitro. To generate phosphorylated Rabs, 4-6mg of purified Rab was incubated with MST3 at a 4:1 - 8:1 wt:wt ratio for 40min to overnight at room temperature in phosphorylation buffer (50 mM Tris-HCl, pH 7.5, 150 mM NaCl, 10 mM MgCl2, 2 mM ATP). FLAG-Rab7A was phosphorylated using TAK1-TAB1 (DU753). 4-6mg of purified Rab7A was incubated with TAK1-TAB1 at a 2:1 wt:wt ratio overnight at room temperature in phosphorylation buffer. Reactions were exchanged into low salt ion exchange buffer (10 mM Tris-HCl, pH 7-8.4 or 10mM MES, pH 5.2, 10 mM NaCl, 1 mM DTT, 5 mM MgCl2) and applied to either a cation (MonoS 5/50 GL) or an anion (MonoQ 5/50 GL) exchanger (GE Healthcare). Phospho FLAG-Rab was eluted with a 0-35% or 0-30% 1M NaCl gradient and confirmed by PhosTag gel electrophoresis, then further purified by gel filtration as above. Rab Ser/Thr →Glu mutations at sites targeted by LRRK2 were generated by PCR site directed mutagenesis using primers ordered from Eurofins Genomics listed in the key resource table. The mutation was confirmed by Sanger sequencing (Eurofins Genomics). The mutant Rabs were expressed in 2xYT and purified as described above. Small scale in vitro kinase assays were performed to indirectly confirm the phospho-site targeted by MST3. 50µg of Rab1A, Rab12, Rab43 or corresponding mutant was incubated with 25µg of MST3 and phosphorylation buffer in a final volume of 75µl overnight at room temperature. 50µg of Rab3A, Rab35 or indicated mutant was incubated with 12.5µg MST3 and phosphorylation buffer in a final volume of 75µl for the indicated time (1 hour – overnight) at room temperature. 50µg of Rab7A or corresponding mutant was incubated with 12.5µg of TAK1-TAB1 (DU753) or MST3 and phosphorylation buffer in a final volume of 75µl overnight at room temperature. All reactions were stopped by the addition of 4X SDS sample buffer, boiled at 96°C for 5 minutes and visualised by 10% Phostag SDS PAGE with Coomassie staining.

Phospho-Rabs for GTP-dependent pulldowns with TBC1D2 were then nucleotide-exchanged. 2-4mg of Rab was incubated with 10mM EDTA (pH 8) and a 5-10 fold excess of GTP or GDP (100mM) in 1ml total volume for 15 minutes at room temperature. MgCl2 was added to 15mM and incubated for a further 15 minutes on ice followed by centrifugation (16,000 x g,10 minutes, 4°C) and gel filtration as previously described.

### Expression and Purification of TBC1D2 constructs

https://dx.doi.org/10.17504/protocols.io.e6nvwx6xdgmk/v1

All wild type and mutant variants of TBC1D2 were expressed and purified as described in protocols.io. The cDNA for all of these constructs of TBC1D2 was ordered from Genscript. This includes Nus-tagged CC region 353–436, which was used for **Fig 6** and for both of the ITC experiments, and the PH domain 43–150 which was used for pulldowns in **Suppl Fig S9.** Further information for these and all other constructs ordered from Genscript can be found in the KRT and in the protocol above. MBP-tagged TBC1D2A-CC (residues 301-555) and Nus-tagged TBC1D2A-N-terminal (residues 1-230) were generated by Ligation Independent Cloning using pLIC-Nus/MBP vectors(Cabrita *et al*, 2006), using primers from Eurofins Genomics. TBC1D2 CC lysine mutants were generated by site-directed mutagenesis using pLIC-MBP-TBC1D2A-CC (301-555) as a template and the indicated primers from Eurofins Genomics. These constructs were expressed in 2xYT as described above.

All TBC1D2 protein variants were purified as described above with the addition of cOmplete™, EDTA-free Protease Inhibitor Cocktail (Merck) during lysis. Following elution from the Ni-agarose, TBC1D2 proteins were further purified by gel filtration chromatography using a Superose 6 Increase 10/300GL column (Cytiva).

### In vitro pulldown with TBC1D2

dx.doi.org/10.17504/protocols.io.dm6gpqk91lzp/v2

The *in vitro* TBC1D2 pull downs were undertaken using techniques described previously in detail. Pulldowns were performed using hexahistidine-tagged TBC1D2 proteins and purified Rabs. Unless otherwise stated, 10µM of the indicated His-tagged TBC1D2 construct was mixed with 10µM of prey phospho-Rab or Rab with 25ul Ni2+-agarose resin in 0.5ml binding buffer (20mM Tris-HCl, pH 8.0, 300mM NaCl, 20mM Imidazole, 5mM MgCl2, 10mM β-mercapotoethanol) and incubated for 15 minutes with gentle shaking. Following centrifugation (1000 x g), the resin was washed three times with 0.5ml binding buffer. Bound proteins were eluted with 50ul elution buffer (20mM Tris-HCl pH 8.0, 300mM NaCl, 200mM imidazole). Samples were visualised with SDS-PAGE and Coomassie Brilliant Blue staining.

### Isothermal titration calorimetry

dx.doi.org/10.17504/protocols.io.j8nlkr7x6v5r/v1

Isothermal titration calorimetry was performed as previously described(Waschbusch *et al*, 2026). Calorimetry experiments were performed twice using an ITC200 with the temperature set at 293K (Particular Sciences). Purified pRab8A was dialyzed together with a fragment of the TBC1D2 CC domain (Nus-tagged 353-436) (20mM Tris-HCl pH8, 150mM NaCl, 5mM MgCl2 and 1mM DTT) for 2-3 hours at 4 °C. The Nus-tag, which contains an N-terminal His-tag, was retained with the CC domain for all steps. However, the Nus-CC domain fusion was subjected to gel filtration chromatography (see purification protocol, above) prior to dialysis with pRab8A. The proteins were then centrifuged at 13,200 x g for 10 minutes prior to injection. 300-400µM of phospho-Rab8 was titrated into 30-40µM of TBC1D2 CC. Data were processed in Origin 7.0 software using a single-site (1:1) binding model (ITC plugin).

### Cultured A549 Cells and generation of A549 cell extract

A549 cells were cultured in 15-cm dishes in DMEM (Gibco) supplemented with 10% (v/v) FBS (Sigma), 2 mM L-glutamine (Gibco), and 100 U/mL penicillin–streptomycin (Gibco), and maintained at 37°C in 5% CO₂. Cells were passaged by detachment with trypsin (Gibco) and split at a ratio of 1:3–1:6. Prior to lysis, cells were treated overnight with 100 nM MLi-2. Cells were washed with PBS (Gibco) and lysed in freshly prepared lysis buffer containing 50 mM Tris-HCl pH 7.4, 150 mM NaCl, 10% (w/v) glycerol, 10 mM sodium β-glycerophosphate, 10 mM sodium pyrophosphate, 0.5% (v/v) NP-40, Complete Mini EDTA-free protease inhibitor (Merck), PhosSTOP phosphatase inhibitor (Merck), and 5 mM MgCl₂. Lysates were clarified by centrifugation at 17,000 × g for 15 min at 4°C, protein concentrations were determined by Bradford assay, and lysates were stored at −80°C.

### Animals and generation of mouse brain extract

C57BL/6J mice were used for this study. Mice were maintained under specific pathogen-free conditions at the University of Dundee (UK). All animal experiments were conducted in accordance with the Animals (Scientific Procedures) Act 1986 and approved by the University of Dundee Animal Welfare and Ethical Review Body under a UK Home Office project licence. Animals were housed under controlled environmental conditions (20–24°C, 45–55% humidity) on a 12 h light/12 h dark cycle with ad libitum access to food and water. Mice received a single subcutaneous injection of vehicle (40% [w/v] 2-hydroxypropyl-β-cyclodextrin; Sigma-Aldrich, #332607) containing MLi-2 LRRK2 inhibitor at a dose of 30 mg kg⁻¹. Two hours following administration, mice were euthanised by cervical dislocation and tissues were rapidly collected, snap-frozen in liquid nitrogen and stored at −80°C until further analysis. Whole brain was rapidly extracted and homogenized in ice cold lysis buffer containing 50 mM Tris-HCl (pH 7.4), 150 mM NaCl, 10% (w/v) glycerol, 10 mM sodium β-glycerophosphate, 10 mM sodium pyrophosphate, 0.5% (v/v) NP-40, Complete Mini EDTA-free protease inhibitor (Merck), PhosSTOP phosphatase inhibitor (Merck), and 5 mM MgCl₂. Lysates were clarified by centrifugation at 17,000 × g for 15 min at 4°C, and protein concentrations were determined using the Bradford assay. Clarified lysates were snap frozen in aliquots and stored until use at −80 °C.

### Immunoprecipitation of pRab and Rab interactors

dx.doi.org/10.17504/protocols.io.5qpvowjn9l4o/v2

The immunoprecipitation of Rab and pRab interactors were performed as described in <u>protocols.io</u>. Phospho-FLAG–Rab (Rab1A, Rab3A, Rab7A, Rab8A, Rab12, Rab35, and Rab43) and FLAG–Rab (Rab1A, Rab3A, Rab7A, Rab8A, Rab12, Rab35, and Rab43) proteins were used as bait for affinity purification experiments to identify differential interacting proteins from cultured A549 cells or mouse brain lysates. Bead volumes and amounts of bait proteins are indicated in the protocol.io deposition. In brief, 20µg of phosphorylated or non-phosphorylated Rab protein pre-conjugated to 5µL of Pierce™ Anti-DYKDDDDK Magnetic Agarose (Thermo Fisher Scientific) by incubation for 1 h at 4°C on a rotating wheel. Following binding, the supernatant was removed and the beads were washed twice with PBS-T (PBS supplemented with 0.01% Tween-20 and 5 mM MgCl₂).A549 cell lysate (400 μg total protein) or mouse brain lysate (750 μg total protein) was then added to the immobilized bait proteins in a final volume of 500 μL and incubated for 1 h at 4°C with rotation. Following incubation, the supernatant was removed and the beads were washed twice with high-salt lysis buffer (lysis buffer supplemented with 500 mM NaCl), followed by a two washes with standard lysis buffer. Beads were subsequently snap-frozen and stored at −80°C prior to preparation for mass spectrometry analysis.

### Sample preparation for quantitative proteomics

https://dx.doi.org/10.17504/protocols.io.8epv5owj6g1b/v1

Sample preparation for quantitative proteomics was performed as previously described in Protocols.io. Immunoprecipitated proteins were eluted from magnetic beads in 40 μL of lysis buffer containing 2% (w/v) SDS, 20 mM HEPES (pH 8.0), Complete EDTA-free protease inhibitor cocktail (Roche), and PhosSTOP phosphatase inhibitor cocktail (Roche). Samples were reduced with 10 mM tris(2-carboxyethyl)phosphine (TCEP) for 30 min at 60°C, followed by alkylation with 40 mM iodoacetamide (IAA) for 30 min at room temperature in the dark.

SDS was then adjusted to a final concentration of 5% (w/v) by addition of 20% SDS, and samples were acidified to a final concentration of 1% (v/v) trifluoroacetic acid (TFA). Proteins were subsequently processed for on-column digestion using micro S-Trap columns (Protifi, USA). Briefly, samples were diluted sixfold with S-Trap binding buffer (90% methanol, 10% 1 M triethylammonium bicarbonate [TEABC]) and loaded onto the S-Trap columns. Columns were centrifuged at 1,000 × g for 1 min, and the flow-through was discarded. Following sample loading, columns were washed four times with 150 μL binding buffer, with centrifugation at 1,000 × g for 1 min after each wash.

On-column digestion was performed by adding 40 μL of 50 mM TEABC (pH 8.0) containing 1 μg Trypsin/Lys-C Mix (MS grade, Promega, UK) and incubating at 47°C for 80 min, followed by overnight incubation at room temperature. Peptides were sequentially eluted with 40 μL 50 mM TEABC (pH 8.0), 40 μL 0.15% (v/v) formic acid, and three successive elutions with 40 μL 80% (v/v) acetonitrile containing 0.15% (v/v) formic acid. Eluted peptides were pooled, dried to completion in a vacuum centrifuge at room temperature, and stored at −20°C until mass spectrometry analysis.

### Quantitative proteomic analysis

dx.doi.org/10.17504/protocols.io.6nvw6pq37gmk/v1

Quantitative proteomic analysis was performed as previously described in Protocols.io. Dried peptides were resuspended in 0.1% (v/v) formic acid containing 0.015% (w/v) n-dodecyl-β-D-maltoside (DDM). Approximately 200 ng of peptide material was analysed using a Vanquish Neo nano-liquid chromatography system coupled online to an Orbitrap Astral mass spectrometer (Thermo Fisher Scientific). Peptides were initially trapped on an Acclaim™ PepMap™ Neo Trap Cartridge (5μm particle size, 300μm diameter, 5mm length) and subsequently separated on an EASY-Spray™ PepMap™ Neo UHPLC column (2μm C18 particle, 150μm diameter, 150mm length). Mobile phase A consisted of 0.1% (v/v) formic acid in water, and mobile phase B consisted of 80% (v/v) acetonitrile containing 0.1% (v/v) formic acid. Peptides were separated using a 13-min gradient with the following conditions: 1% B from 0–0.7 min at 1.8 μL/min; 4% B from 0.7–1.0 min at 1.8 μL/min; 8% B from 1.0–7.7 min at 1.8 μL/min; 22.5% B from 7.7–11.4 min at 1.8 μL/min; 35% B from 11.4–11.8 min at 1.8 μL/min; 55% B from 11.8–12.3 min at 2.5 μL/min; and 99% B from 12.3–13.0 min at 2.5 μL/min. Mass spectrometric analysis was performed in data-independent acquisition (DIA) mode. Full MS (MS1) scans were acquired at a resolution of 240,000 over an m/z range of 380–980, using a normalized AGC target of 500% and a maximum injection time of 3 ms. DIA scans were acquired across the precursor mass range using 4 m/z isolation windows, with fragment ion spectra collected over an m/z range of 150–2,000 and a normalized collision energy (NCE) of 25%. Raw data were processed using DIA-NN software as described below.

### Raw mass spectrometry data analysis

Raw mass spectrometry data were analysed as previously described in Protocols.io. DIA data were processed using DIA-NN (version 1.9.2) with in silico–predicted spectral libraries generated from the reviewed UniProt reference proteomes for either human (A549 cell lysates) or mouse (brain lysates). The UniProt databases were downloaded in January 2023 and contained 20,381 human protein entries plus isoforms and 17,124 mouse protein entries plus isoforms. Database searches were performed using strict trypsin specificity, allowing cleavage C-terminal to lysine and arginine residues, with a maximum of one missed cleavage permitted per peptide. Peptides ranging from 7 to 30 amino acids in length were considered for identification using DIA-NN default settings. Carbamidomethylation of cysteine residues was specified as a fixed modification. Protein and peptide identifications were filtered using the default DIA-NN false discovery rate (FDR) settings, and quantitative analyses were performed using the resulting protein group intensity values. Subsequent statistical analyses and data visualisation were carried out as described below. The code for processing of mass spectrometry data is provided at the following GitHub link: https://github.com/M-Jaconelli/JupyterNB-MassSpectrometry-DataAnalysis.

### Statistical Analysis and Data Visualization

DIA-NN output files were processed and visualised using Python (version 3.11.5; Project Jupyter) with the packages pandas (v2.0.3), NumPy (v1.26.1), Matplotlib (v3.7.2), and Seaborn (v0.12.2). Processed datasets were subsequently uploaded to the CURTAIN web application for interactive exploration and visualisation. Proteins identified by a single peptide were excluded from further analysis. Protein group intensities were log₂-transformed and filtered to retain proteins detected in 100% of replicates within at least one experimental condition. Missing values were imputed by random sampling from a Gaussian distribution centred on the 1st percentile of the global protein intensity distribution for each sample. Differential abundance analysis was performed using the Limma package (version 3.56.2) in R (version 4.3.1). Linear models were fitted to the data using the lmFit function, and empirical Bayes moderation was applied using the eBayes function to improve variance estimation. Statistical significance was determined using moderated t-statistics, with multiple testing corrections performed using the Benjamini–Hochberg procedure to control the false discovery rate (FDR) at 5%. Data visualisation included volcano plots, scatter plots, heatmaps, and protein abundance comparisons generated using Python and CURTAIN. Proteins with an adjusted P value (FDR) < 0.05 were considered statistically significant unless otherwise stated.

### Data search portal and visualisation

To make the data available for anyone to query and search for any protein of interests within the generated datasets, we had imported the analysed data and provided a search portal for them through Cinder. Cinder was created using Django v5 (https://www.djangoproject.com/) as backend (https://github.com/noatgnu/cinderbackend) and Angular Web Framework v22 (https://angular.dev/) as frontend (https://github.com/noatgnu/CinderFrontend) with persistent database based on Postgres v14 (https://www.postgresql.org/). The search output and differential analysis were ingested by Cinder where they are broken into multiple segments due to maximum length of text document that might be stored within the database itself. When a user search using one or multiple search terms, each term would be quickly search through all text segments belonging to all documents of the data collection. Then further processing would be done to identify which row exactly in each table that the term of interests was found. Results were then filtered depending on whether the user looking for a gene name, UniProt accession id, or experiment specific unique identifier.

In the frontend, for each result from the collection portal, the intensity across different samples with the detected protein would be shown in average bar chart for each condition. For the differential analysis, we also provide heatmap plot depicting the direction of changes based on the log2fc of the comparison. All the plots provided above were done using plotly.js (https://plotly.com/javascript/). Cinder data share portal also provides multiple options for the user as well as the collection creators to further customize like the size, color, order of conditions, significant filter, and more.

### Confirmation of *in vitro* pRab phosphorylation sites with mass spectrometry

<u>dx.doi.org/10.17504/protocols.io.6wgq9rqwxlk5/v1</u>

Confirmation of *in vitro* Rab phosphorylation sites by mass spectrometry was performed as described in Protocols.io. For each Rab protein (RAB1A, RAB3A, RAB7, RAB8A, RAB12, RAB35, and RAB43), 1 μg of purified phosphorylated FLAG–Rab protein was diluted 1:39 in 50 mM triethylammonium bicarbonate (TEABC; pH 8.0). Samples were reduced with 10 mM tris(2-carboxyethyl)phosphine (TCEP) at 60°C for 30 min with agitation at 1,100 rpm using a ThermoMixer C (Eppendorf). Cysteine residues were subsequently alkylated with 40 mM iodoacetamide (IAA) for 30 min at room temperature with agitation at 1,100 rpm in the dark.

Proteins were digested overnight (∼16 h) at 37°C following the addition of 100 ng Trypsin/Lys-C Mix, MS grade (Thermo Fisher Scientific). Digestion was terminated by acidification with formic acid to a final concentration of 0.1% (v/v). Peptides were desalted using C18 EvoTips (Evosep), eluted with 40% (v/v) acetonitrile, dried by vacuum centrifugation, and stored at −20°C until analysis.

For LC–MS/MS analysis, peptides were resuspended in 0.1% (v/v) formic acid containing 0.015% (w/v) n-dodecyl-β-D-maltoside (DDM). Approximately 5 ng of peptide material was analysed using an Ultimate 3000 UHPLC system coupled online to an Orbitrap Fusion Lumos Tribrid mass spectrometer (Thermo Fisher Scientific). Peptides were separated on an EASY-Spray analytical column (75 μm × 50 cm, 2 μm C18 particles) equipped with an integrated nano-electrospray emitter. Mobile phase A consisted of 0.1% (v/v) formic acid in water, and mobile phase B consisted of 80% (v/v) acetonitrile containing 0.08% (v/v) formic acid.

Peptides were separated using a 60-min gradient at a flow rate of 0.3 μL/min as follows: 3% B at 0 min; 5% B at 5 min; 7% B at 7 min; 35% B at 40 min; 95% B at 50–54 min; and re-equilibration at 3% B from 55–60 min.

Detailed instrument acquisition parameters are available in the deposited raw data files within the PRIDE repository (see Data Availability). Briefly, samples were analysed using a semi-targeted data-dependent acquisition (ddMSn) workflow. Full MS scans were acquired in the Orbitrap at a resolution of 120,000 over an *m/z* range of 375–1,500, with a normalized AGC target of 100% and a maximum injection time of 50 ms. Data-dependent MS/MS scans were acquired from precursor ions with charge states of 2–7 using a Top10 method and an isolation window of 1.6 *m/z*. Electron-transfer/higher-energy collision dissociation (EThcD) fragmentation was employed throughout the analysis.

A targeted inclusion list comprising the expected phosphopeptide sequences from all phosphorylated Rab proteins was used to prioritize acquisition of relevant spectra. When no inclusion-list targets were detected, conventional data-dependent acquisition was performed. MS/MS spectra were acquired in the Orbitrap at a resolution of 30,000 using a normalized AGC target of 100% and a maximum injection time of 120 ms. Raw data files were processed using Proteome Discoverer version 2.4 (Thermo Fisher Scientific) with Mascot version 2.6.2 (Matrix Science) as the search engine. Spectra were searched against an in-house protein sequence database (MRC database) using a precursor mass tolerance of 10 ppm and a fragment mass tolerance of 0.06 Da. Searches were performed with strict trypsin specificity, allowing up to two missed cleavages. Phosphorylation of serine, threonine, and tyrosine residues was specified as a variable modification, while carbamidomethylation of cysteine residues was set as a fixed modification.

All phosphopeptide identifications were manually inspected to confirm phosphorylation site localisation. Confidence in site assignment was assessed using ptmRS localisation probabilities, DeltaM mass accuracy values, spectral quality, and the number of supporting peptide-spectrum matches.

### Consurf evolutionary analysis and AlphaFold modelling

ConSurf sequence analyses was performed with 405 homologous protein sequences from 394 species obtained from the mammalian dataset in OrthoDB(v13.0). (Yariv *et al*., 2023). Sequences were supplied in FASTA format and aligned using MAFFT, with the evolutionary model setting set to BEST through Contort (https://contort.proteo.nexus/). Contort is an alternative user interface for running ConSurf analysis (https://github.com/Rostlab/ConSurf). It was created to run analyses through a user interface on personal hardware without complex command-line execution. This work leverages a design framework similar to Cinder. The source code available at https://github.com/noatgnu/Contort (frontend),https://github.com/noatgnu/contort-backend (backend), and https://github.com/noatgnu/ConSurf-StandAlone/tree/1.04 (the copy of the command-line ConSurf version that we built upon). In order to perform 3-D modelling of the interaction between TBC1D2 and Rab8A, we used Rab8A phosphorylated on Thr72, the physiological LRRK2 phosphorylation site and predicted its complex with TBC1D2 using AlphaFold3 (Abramson *et al*., 2024). Confidence models suggested that phosphorylated Rab8A binds to a dimeric form of TBC1D2 through its coiled-coil domain, generating a heterotetrameric complex reminiscent of numerous previously characterised Rab–effector assemblies (Khan & Menetrey, 2013). The outputs including the confidence statistics for the modelling, have been deposited in Zenodo (see below, **Data Availability**).

## Data availability

The mass spectrometry proteomics data generated in this study have been deposited with the ProteomeXchange Consortium via the PRIDE partner repository under dataset identifier PXD084766. The code used for data processing and analysis are available through GitHub (github.com/M-Jaconelli/JupyterNB-MassSpectrometry-DataAnalysis).

Experimental protocols for DIA mass spectrometry data analysis have been deposited (dx.doi.org/10.17504/protocols.io.6nvw6pq37gmk/v1). The Rab3A/pRab3A mouse brain interactome dataset analysed in this study was generated as part of a previously published study (PRIDE identifier PXD065125). The data have been re-analysed here alongside newly generated datasets using the same bioinformatic and statistical workflow. The original data remain available through the repositories associated with the previous publication. CURTAIN links to the proteomics data are available in Excel format in **Suppl Table 1**. ITC data from experiments performed on the Microcal ITC200 instrument are available in Zenodo as Origin 7.0 format files (10.5281/zenodo.23820352). The full output of sequence conservation from Consurf shown in **Fig 5C** has been deposited (10.5281/zenodo.23848038). Uncropped immunoblots corresponding to **Fig 5D**, and associated graphing data in **Fig 5E** (GraphPad Prism, raw Excel data) are deposited in the Zenodo repository under 10.5281/zenodo.23891932. Raw gels for **Suppl Fig S1** have been deposited in Zenodo under 10.5281/zenodo.23930544. Data associated with the AlphFold prediction and quality metrics shown in **Fig 5B** and **Suppl Fig S8** have been deposited in Zenodo under 10.5281/zenodo.23860938. Raw SDS-PAGE gels used to generate **Fig 6** and **Suppl Fig S9** have been deposited in the Zenodo server (10.5281/zenodo.23931555). Raw SDS-PAGE and PhosTag gel for **Fig 1**, as well as the raw immunoblot used to generate **Suppl Fig S2**, have also been deposited in the Zenodo server (10.5281/zenodo.23896933).

## ACKNOWLEDGMENTS

We thank the excellent technical support of the MRC-PPU DNA sequencing service (coordinated by Gary Hunter), the MRC-PPU tissue culture team (coordinated by Dr Edwin Allen), the MRC-PPU mass spectrometry facility team (coordinated by Dr Renata Soares), and the MRC-PPU Reagents and Services team (coordinated by Dr. James Hastie). A.R.K. was funded by Research Ireland through a Research Frontiers grant, 20/FFP-A/8446. This work was supported by the UK Medical Research Council (MC_UU_00018/1) (D.R.A.) and also supported as part of Aligning Science Across Parkinson’s (ASAP) (https://ror.org/03zj4c476) Team Alessi ASAP[000463] and Team Pfeffer ASAP[027054], a research initiative managed by the Coalition for Aligning Science (CAS) (https://ror.org/00rf4ez40). The Michael J. Fox Foundation for Parkinson’s Research (MJFF) (https://ror.org/03arq3225) is the funding and “strategic partner.” For the purpose of open access, the author has applied a CC BY public copyright license to all Author Accepted Manuscripts arising from this submission. We thank Suzanne Pfeffer for helpful advice and discussions.

## AUTHOR CONTRIBUTIONS

A.R.K. and D.R.A. conceived, designed and supervised the study. E.N. expressed and purified recombinant proteins, performed in vitro pulldowns and calorimetry. E.N. prepared FLAG-tagged pulldowns of proteins from cell lysates/brain in preparation for mass spectrometry. J.S. expressed proteins in HEK293 cells, and performed co-IPs, AlphaFold and ConSurf analysis. M.J. performed all mass spectrometry experiments and analyses. T.P. generated the tool to compare candidate Rab effector enrichment across phosphorylated and non-phosphorylated Rabs in A549 and brain extracts. All authors contributed to writing of this manuscript.

## Conflicts of Interest

The authors declare no conflicts of interest.

## Supplemental Figures

**Suppl Fig S1:** Comparisons of the migration of FLAG-Rab species following *in vitro* kinase phosphorylation of WT and Ser/Thr→Glu mutations. All of these analyses were performed at small scale to analyze efficiency of phosphorylations under various conditions (**see Materials and Methods**). In all cases, the red dotted rectangles indicate successful phosphorylation, which occurs only with the addition of the WT protein, the kinase and ATP. MST3 was used for all Rabs, except for Rab7A which was phosphorylated more efficiently by TAK1 kinase. **(A)** Phosphorylation of Rab1A WT and T75E, showing successful reaction with WT protein. **(B)** Phosphorylation of Rab3A at various time points indicated. Optimal times of incubation are 1-4 hours. **(C)** Phosphorylation of Rab7A, which migrates much slower compared to other phospho-Rabs in PhosTag gels. The S72E variant appears as a doublet for the non-phosphorylated species. TAK1 is significantly better than MST3 at phosphorylating Rab7A. **(D)** Phosphorylation of Rab12, which appears as a diffuse band in PhosTag gels. **(E)** Phosphorylation of Rab35 at various time points. **(F)** Phosphorylation of Rab43, revealing successful reaction with WT but not the T82E mutation.

**Suppl Fig S2**: Cell lysate pulldowns and IP confirm known interactions between phospho-Rab8A and RILP proteins with optimised lysis and wash conditions. FLAG-pRab8A and FLAG-Rab8A were immobilised on anti-FLAG magnetic beads and incubated with 300 μg A549 cell lysate in duplicate. Beads were washed with high salt lysis buffer (HS) followed by low salt lysis buffer (LS) or low salt lysis buffer alone, bound proteins were eluted and visualised by SDS PAGE. Membranes were cut and sections probed separately for LRRK2, RILPL1 and RILPL2. Controls include beads and lysate incubated together without Rab bait, FLAG-pRab8A and FLAG-Rab8A incubated with beads without lysate, and beads alone to confirm the specificity of the bands detected. RILPL1 and RILPL2 were enriched in pRab8A pulldowns compared to Rab8a confirming the specificity of the IP. A 20μg lysate input is shown as a loading reference. The stringent conditions in these trials involving 2 HS and 2 LS washes was used for subsequent MS identification (see **Materials and Methods**).

**Suppl Fig S3:** Differential interactome of phosphorylated vs non-phosphorylated Rab1A. Volcano plots show proteins from A549 cell extract and mouse brain lysate. STK24(MST3) is denoted with an asterisk (*) since it is a contaminant from the kinase reaction that generated pRab1A.

**Suppl Fig S4:** Differential interactome of phosphorylated vs non-phosphorylated Rab3A. The volcano plots show proteins from A549 cell extract. The corresponding interactome from mouse brain lysates has previously been reported(Chen *et al*., 2026). STK24(MST3) is denoted with an asterisk (*) since it is a contaminant from the kinase reaction that generated pRab3A.

**Suppl Fig S5:** Differential interactome of phosphorylated vs non-phosphorylated Rab7A. Volcano plots show proteins from A549 cell extract and mouse brain lysate. TAK1/TAB1 are indicated with asterisks (*) since they are contaminants from the kinase reaction that generated pRab7A.

**Suppl Fig S6:** Differential interactome of phosphorylated vs non-phosphorylated Rab12. Volcano plots show proteins from A549 cell extract and mouse brain lysate.

**Suppl Fig S7:** Differential interactome of phosphorylated vs non-phosphorylated Rab35. Volcano plots show proteins from A549 cell extract and mouse brain lysate.

**Suppl Fig S8:** Modeling and prediction quality of TBC1D2 CC domain and its complex with pRab8A. **(A)** Structural modelling was performed using AlphaFold3(Abramson *et al*., 2024) to investigate the interaction between phosphorylated Rab8A and the coiled-coil domain of TBC1D2. Predictions included two molecules of Rab8A, with Thr72 specified as a phosphorylated residue and GTP and Mg²⁺ ions included in the input, together with two copies of the TBC1D2 coiled-coil domain (residues 300-555). Five models were generated, all predicting a similar heterotetrameric arrangement comprising two molecules of phospho-Rab8A and a dimer of TBC1D2 coiled-coil domains. The highest-ranked model (ipTM = 0.53; pTM = 0.56) was selected for visualisation, with a view highlighting the predicted interaction interface involving TBC1D2 residues 402-435. Structural confidence was assessed using per-residue predicted local distance difference test (pLDDT) scores, represented by a colour scale ranging from dark blue (very high confidence) through light blue (high confidence) and yellow (low confidence) to orange (very low confidence). Residue K410 is labeled on each of the 2 chains of the coiled coil dimer (K414 is not as prominent in this view). **(B)** Predicted aligned error (PAE) plots for the complete heterotetrameric complex were generated using PAE Viewer (https://pae-viewer.uni-goettingen.de/), with red rectangles highlighting the intermolecular interfaces of interest.

**Suppl Fig S9:** *In vitro* pulldowns confirm cellular co-IPs and reveal that both lysines in the CC domain of TBC1D2 are required for complex formation with pRab8A. **(A)** Purified proteins used for these experiments are shown (left) with His-tagged TBC1D2 as bait. The variants of TBC1D2 are MBP-tagged 301-555 (CC domain), Nus-tagged 1-230 N-terminal region (Nter), or the core PH domain 43-150. The prey (untagged proteins) in these pulldowns were (p)Rab8A as indicated. Bait and prey proteins were 10μM final concentration. **(B)** His-tagged WT and mutants of MBP-tagged TBC1D2 (301-555, CC domain) were used as bait to pulldown pRab8A. The mutants were K410E, K414E, K410E/K414E, and K410A/K414A. None of these mutants were able to interact with pRab8A. Protein concentrations for bait and prey proteins were 5μM. **(C)** *In vitro* pulldown assays of phosphorylated Rabs using His-tagged TBC1D2 CC domain (301-555, MBP-tagged). Only pRab3A was pulled down with the CC domain of TBC1D2 – pRab7A and pRab43 did not interact under the same conditions. Protein concentrations for the pulldowns were set at 5μM.

## Notes

https://cinder.proteo.info/#/collate/view/11

https://curtain.proteo.info/#/1e64d1dc-de70-4217-9a8f-6adad186ce93

https://curtain.proteo.info/#/9d675777-929b-4f7c-9bf5-985a63394741

https://curtain.proteo.info/#/73365b43-7560-4da1-a641-952f6ac9c4cc

https://curtain.proteo.info/#/6b02c724-db32-4794-84e3-678c0752e11d

https://curtain.proteo.info/#/68c38f1b-4016-4dd7-b13b-59d925ff09ee

https://curtain.proteo.info/#/9a4fd6bf-a2fd-4579-b6ad-206b2983ca6b

https://curtain.proteo.info/#/47e1361a-6a83-41b4-b8e5-6aa3ca3c1527

https://curtain.proteo.info/#/3c2e09a1-3e22-4548-9d6f-28379dff6951

https://curtain.proteo.info/#/330e114b-3f10-44d7-9a68-227bdc5771b9

### Competing Interest Statement

The authors have declared no competing interest.

