## Supplementary material for "Defining phospho-Rab signalling networks downstream of LRRK2 kinase": Suppl Fig S1

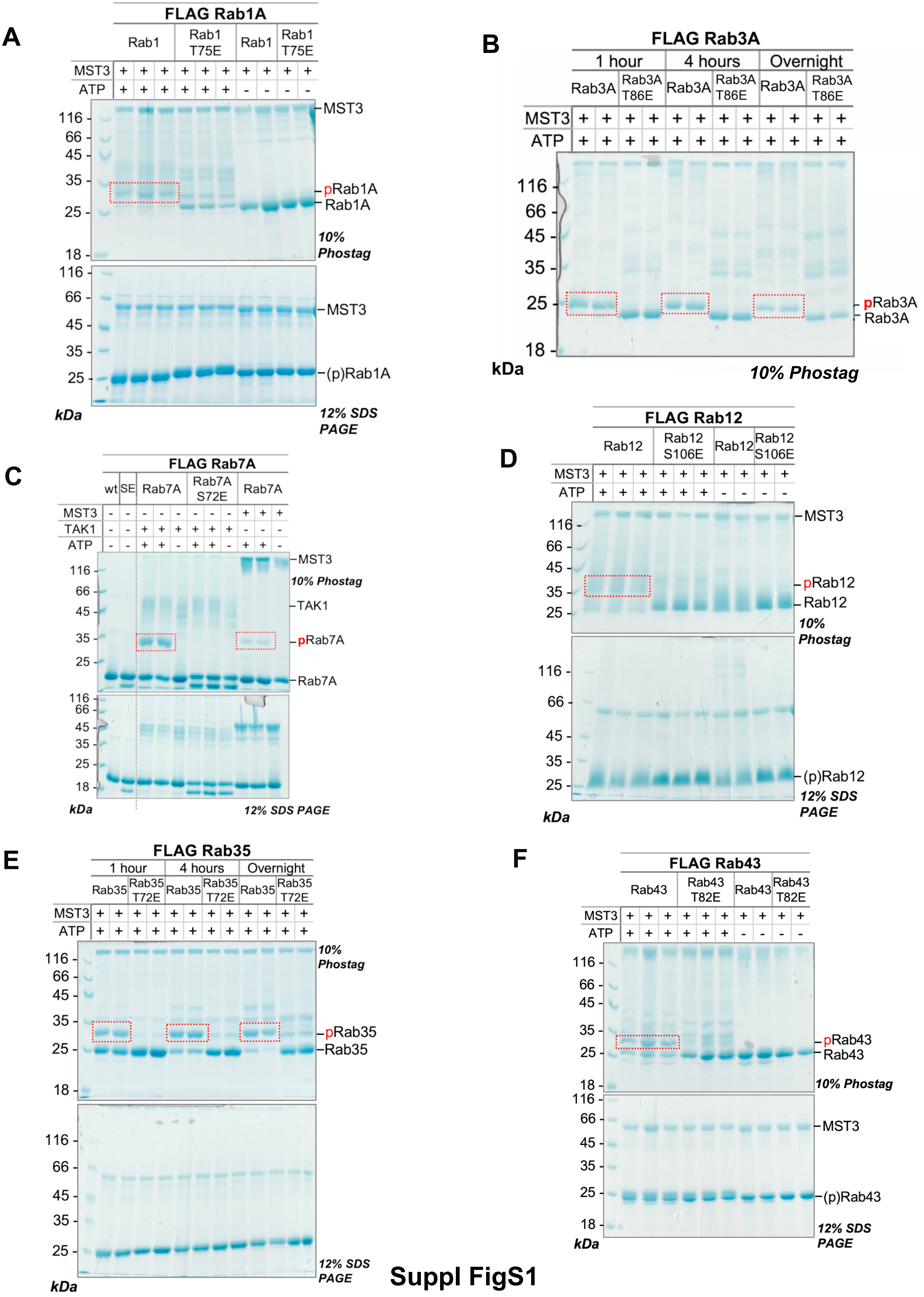

A
FLAG Rab1A
Rab1
Rab1 T75E
Rab1
Rab1 T75E
MST3
ATP
116
66
45
35
25
18
MST3
pRab1A
Rab1A
10% Phostag
(p)Rab1A
kDa
12% SDS PAGE
B
FLAG Rab3A
1 hour
4 hours
Overnight
Rab3A
Rab3A T86E
pRab3A
10% Phostag
C
FLAG Rab7A
wt
SE
Rab7A
Rab7A S72E
TAK1
pRab7A
Rab7A
D
FLAG Rab12
Rab12
Rab12 S106E
pRab12
(p)Rab12
E
FLAG Rab35
Rab35
Rab35 T72E
pRab35
F
FLAG Rab43
Rab43
Rab43 T82E
pRab43
(p)Rab43


Suppl FigS1
