## Supplementary material for "Defining phospho-Rab signalling networks downstream of LRRK2 kinase": Suppl Fig S2

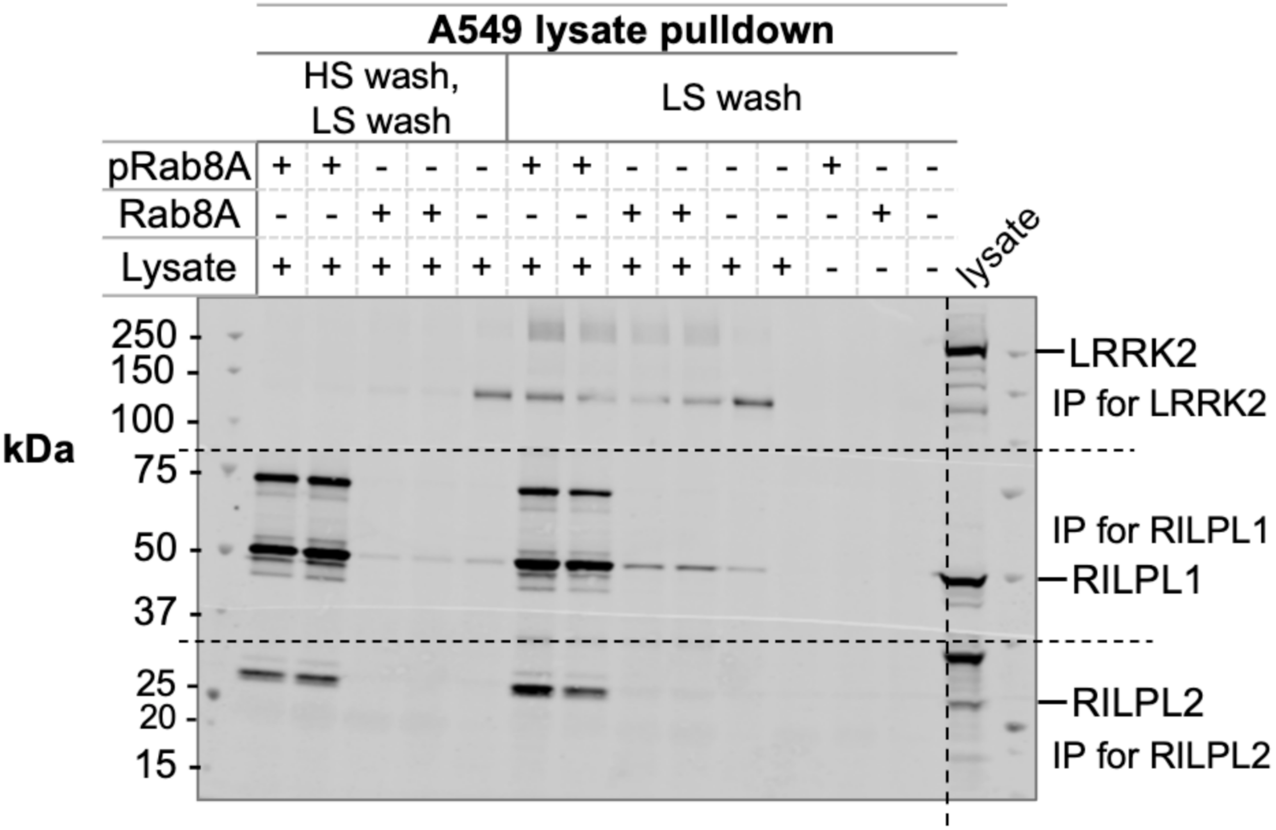
A549 lysate pulldown
HS wash, LS wash
LS wash
pRab8A
Rab8A
Lysate
lysate
kDa
250
150
100
75
50
37
25
20
15
LRRK2
IP for LRRK2
IP for RILPL1
RILPL1
RILPL2
IP for RILPL2

**Suppl FigS2**
