## Supplementary material for "Defining phospho-Rab signalling networks downstream of LRRK2 kinase": Suppl Fig S3

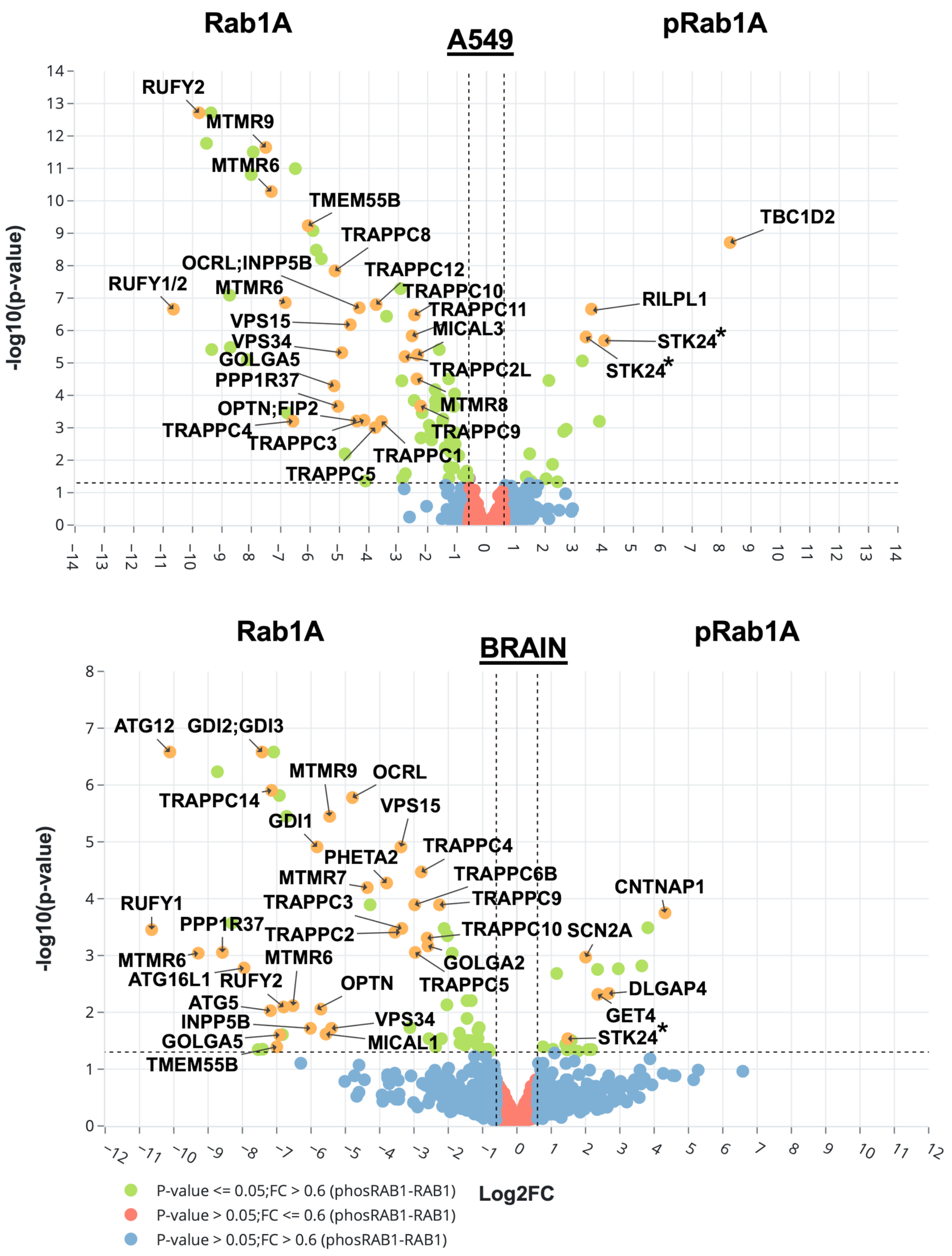
Rab1A
A549
pRab1A
-log10(p-value)
RUFY2
MTMR9
MTMR6
TMEM55B
TRAPPC8
OCRL;INPP5B
TRAPPC12
RUFY1/2
MTMR6
TRAPPC10
TRAPPC11
VPS15
MICAL3
VPS34
GOLGA5
TRAPPC2L
PPP1R37
MTMR8
OPTN;FIP2
TRAPPC4
TRAPPC9
TRAPPC3
TRAPPC1
TRAPPC5
TBC1D2
RILPL1
STK24*
STK24*
Rab1A
BRAIN
pRab1A
-log10(p-value)
ATG12
GDI2;GDI3
MTMR9
OCRL
TRAPPC14
VPS15
GDI1
TRAPPC4
PHETA2
MTMR7
TRAPPC6B
TRAPPC3
TRAPPC9
CNTNAP1
RUFY1
PPP1R37
TRAPPC2
TRAPPC10
SCN2A
MTMR6
MTMR6
GOLGA2
ATG16L1
RUFY2
OPTN
TRAPPC5
DLGAP4
ATG5
GET4
INPP5B
VPS34
STK24*
GOLGA5
MICAL1
TMEM55B
Log2FC
P-value <= 0.05;FC > 0.6 (phosRAB1-RAB1)
P-value > 0.05;FC <= 0.6 (phosRAB1-RAB1)
P-value > 0.05;FC > 0.6 (phosRAB1-RAB1)

Suppl Fig S3
