## Supplementary material for "Defining phospho-Rab signalling networks downstream of LRRK2 kinase": Suppl Fig S4

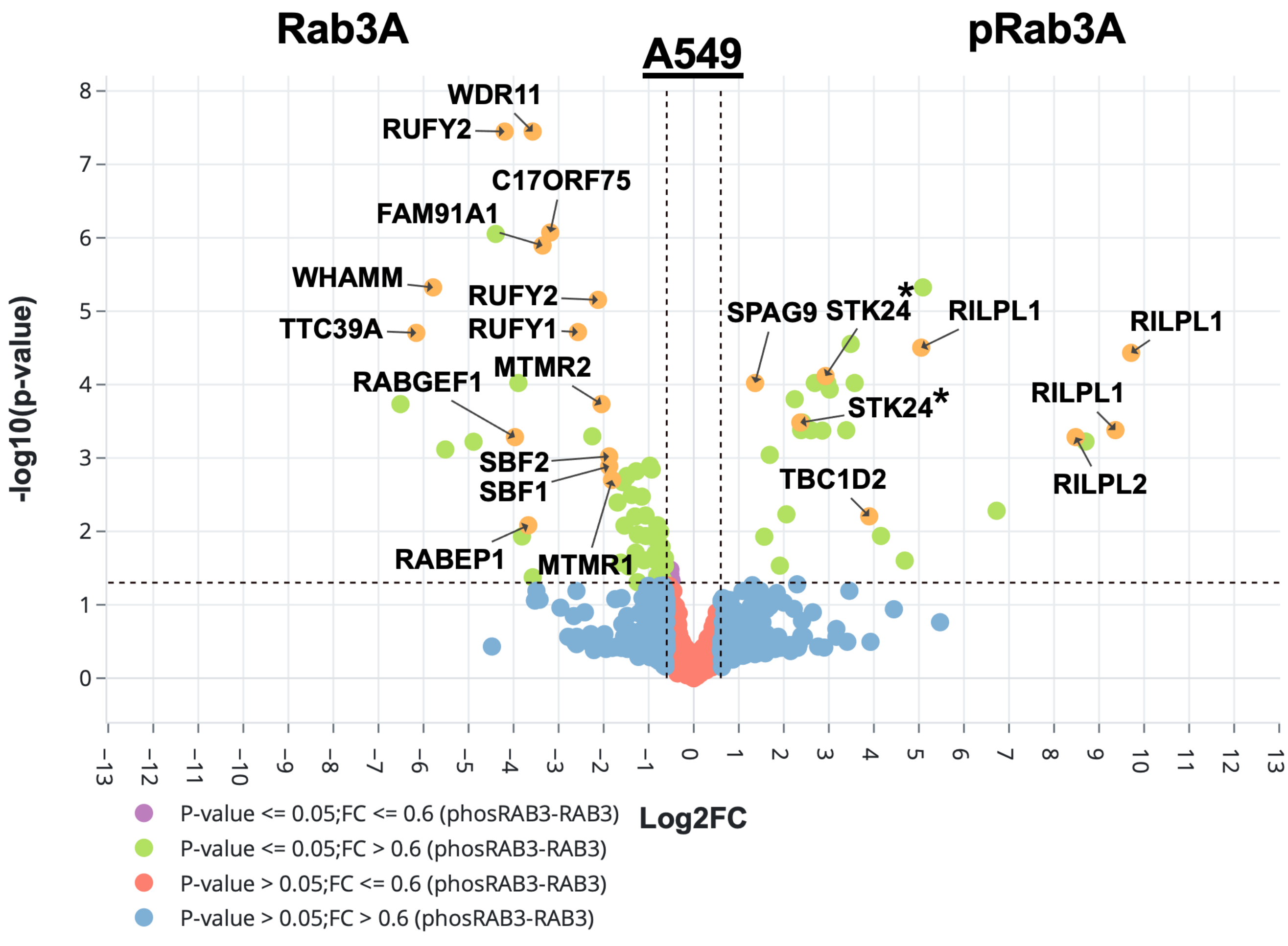

Rab3A
A549
pRab3A
WDR11
RUFY2
C17ORF75
FAM91A1
WHAMM
RUFY2
TTC39A
RUFY1
MTMR2
RABGEF1
SBF2
SBF1
RABEP1
MTMR1
SPAG9
STK24*
RILPL1
RILPL1
STK24*
RILPL1
RILPL2
TBC1D2
-log10(p-value)
Log2FC
P-value <= 0.05;FC <= 0.6 (phosRAB3-RAB3)
P-value <= 0.05;FC > 0.6 (phosRAB3-RAB3)
P-value > 0.05;FC <= 0.6 (phosRAB3-RAB3)
P-value > 0.05;FC > 0.6 (phosRAB3-RAB3)


**Suppl Fig S4**
