## Supplementary material for "Defining phospho-Rab signalling networks downstream of LRRK2 kinase": Suppl Fig S5

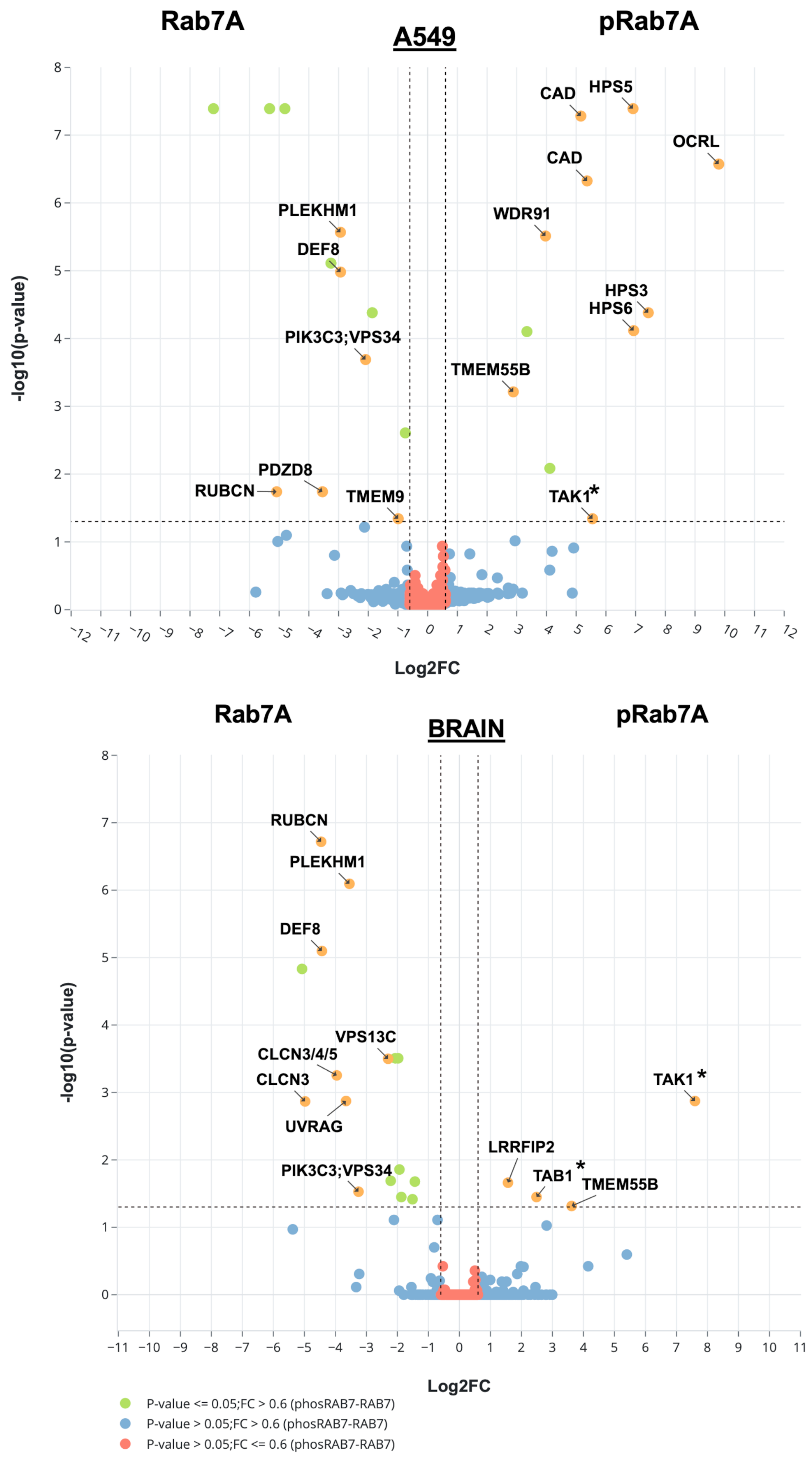
Rab7A
A549
pRab7A
-log10(p-value)
Log2FC
CAD
HPS5
OCRL
CAD
PLEKHM1
WDR91
DEF8
HPS3
HPS6
PIK3C3;VPS34
TMEM55B
PDZD8
RUBCN
TMEM9
TAK1*
Rab7A
BRAIN
pRab7A
-log10(p-value)
Log2FC
RUBCN
PLEKHM1
DEF8
VPS13C
CLCN3/4/5
CLCN3
UVRAG
TAK1*
LRRFIP2
PIK3C3;VPS34
TAB1*
TMEM55B
P-value <= 0.05;FC > 0.6 (phosRAB7-RAB7)
P-value > 0.05;FC > 0.6 (phosRAB7-RAB7)
P-value > 0.05;FC <= 0.6 (phosRAB7-RAB7)

**Suppl Fig S5**
