## Supplementary figures and images for "Defining phospho-Rab signalling networks downstream of LRRK2 kinase"

### Suppl Fig S6

A

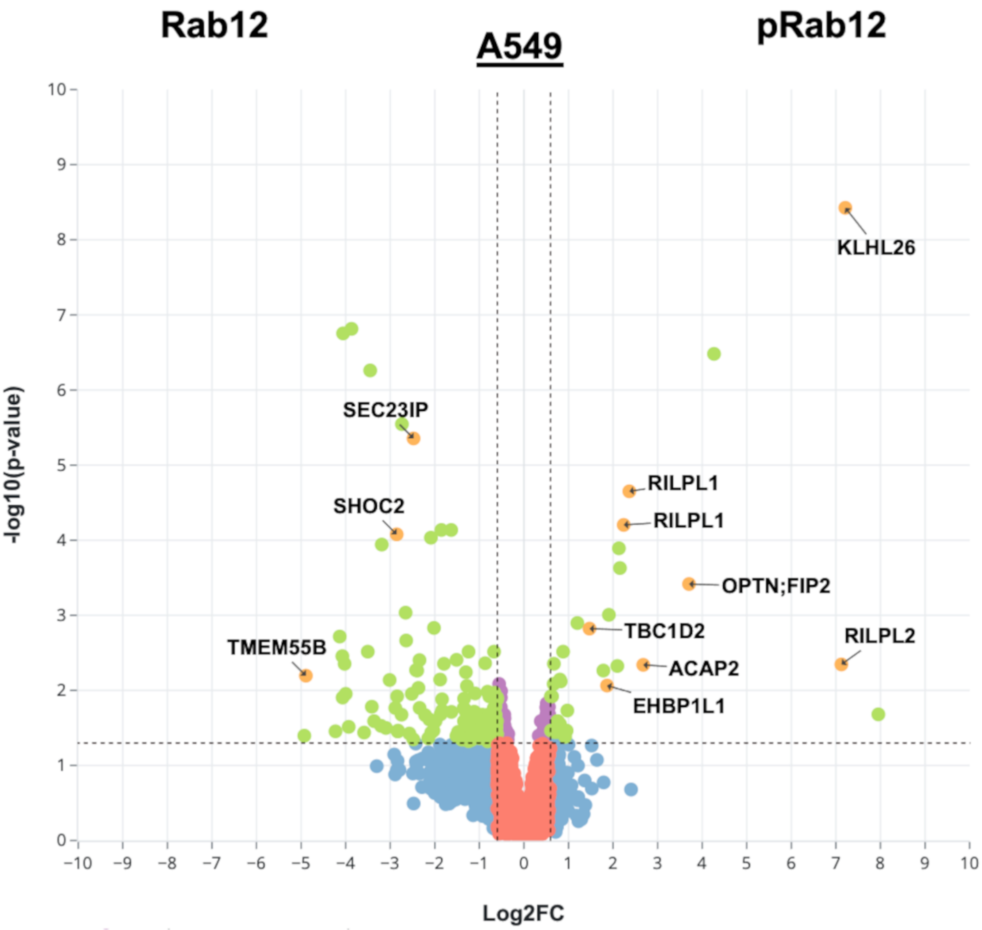


B

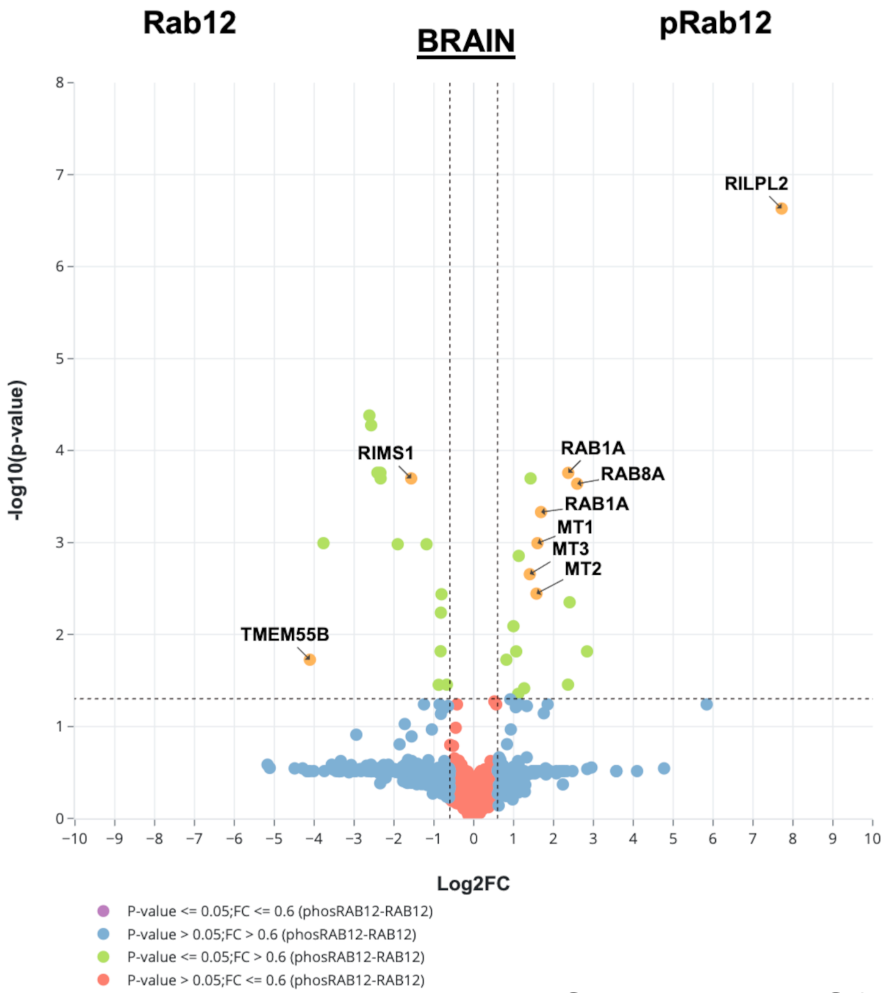


**Suppl Figure S6**

### Suppl Fig S7

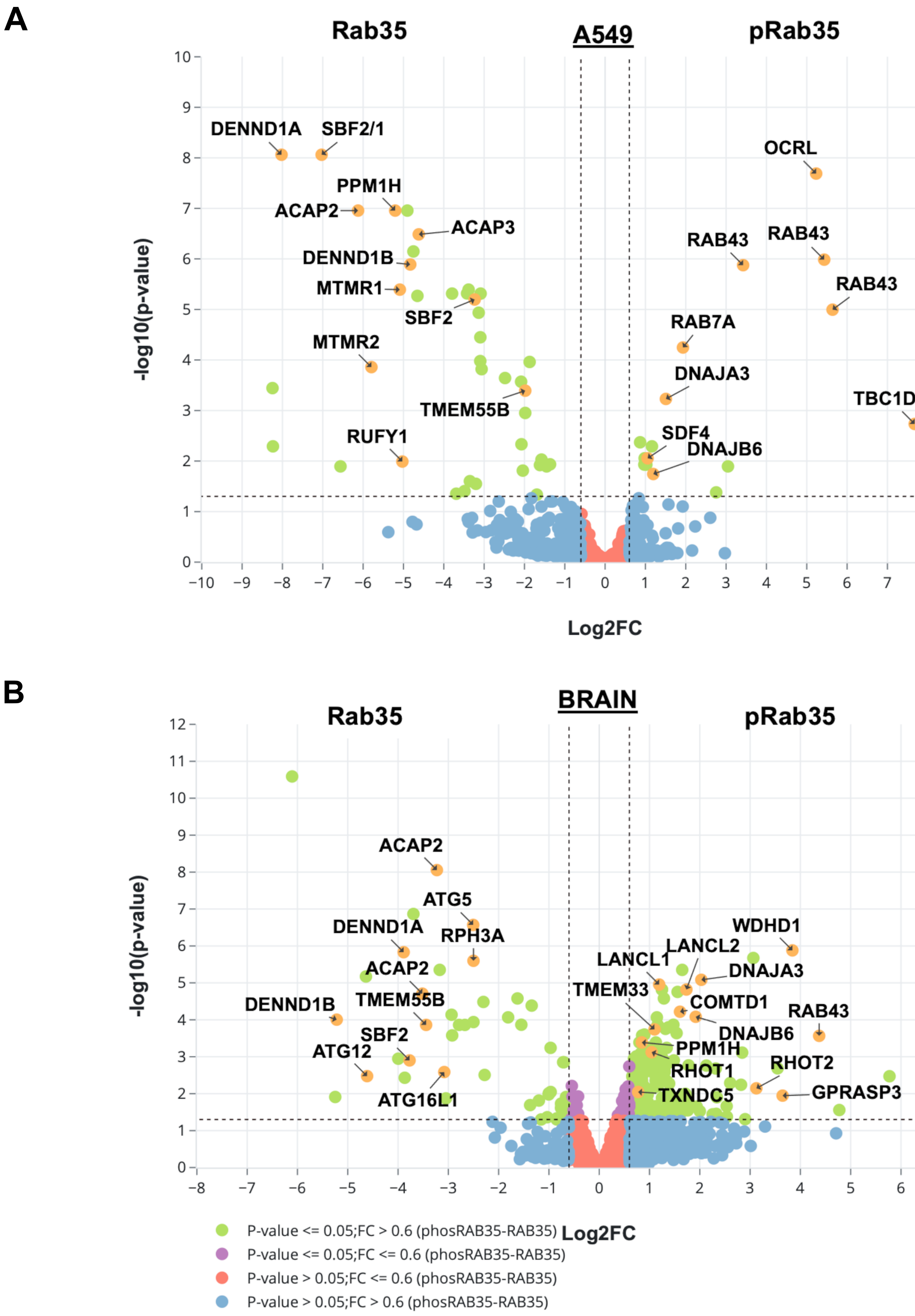


**Suppl Figure S7**

### Suppl Fig S8

A

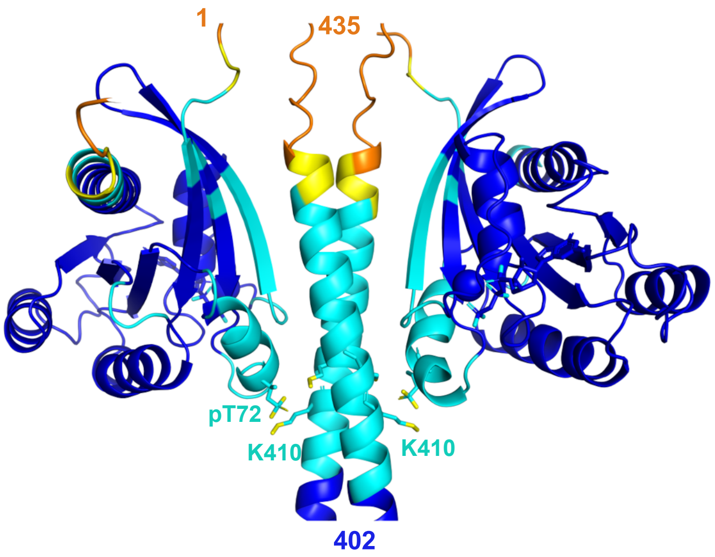


B

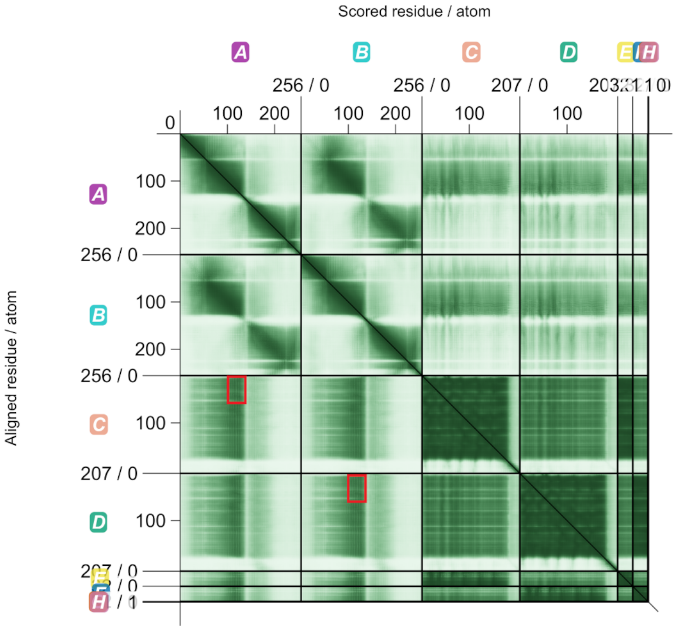


**Suppl Fig S8**
