## Supplementary material for "Defining phospho-Rab signalling networks downstream of LRRK2 kinase": Suppl Fig S9

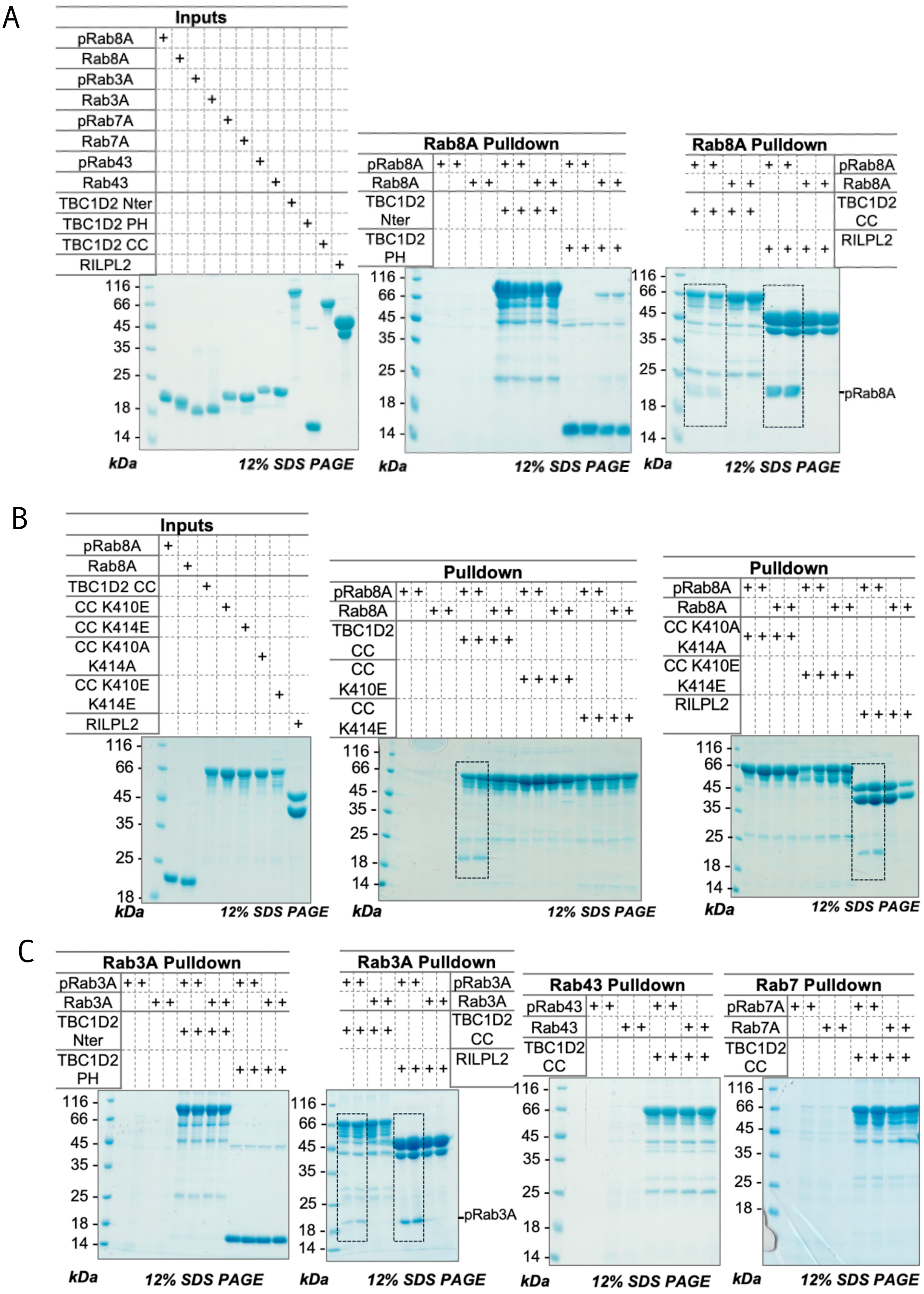
A
Inputs
pRab8A
Rab8A
pRab3A
Rab3A
pRab7A
Rab7A
pRab43
Rab43
TBC1D2 Nter
TBC1D2 PH
TBC1D2 CC
RILPL2
Rab8A Pulldown
TBC1D2 Nter
TBC1D2 PH
TBC1D2 CC
RILPL2
pRab8A
kDa
12% SDS PAGE
B
Inputs
TBC1D2 CC
CC K410E
CC K414E
CC K410A K414A
CC K410E K414E
RILPL2
Pulldown
C
Rab3A Pulldown
pRab3A
Rab43 Pulldown
pRab43
Rab43
Rab7 Pulldown
pRab7A
Rab7A
116
66
45
35
25
18
14

**Suppl Fig S9**
