## Supplementary material for "Defining phospho-Rab signalling networks downstream of LRRK2 kinase": Suppl Table 1

### Supplementary Table 1: CURTAIN links to volcano plots of mass spectrometry data of phospho-Rab interactors

| **Source** | **RAB IP** | **Curtain Link** | **-P** | **+P** |
|---|---|---|---|---|
| A549 | 1 | https://curtain.proteo.info/#/1e64d1dc-de70-4217-9a8f-6adad186ce93 | 73 | 14 |
| A549 | 3 | https://curtain.proteo.info/#/9d675777-929b-4f7c-9bf5-985a63394741 | 50 | 25 |
| A549 | 7 | https://curtain.proteo.info/#/73365b43-7560-4da1-a641-952f6ac9c4cc | 12 | 11 |
| A549 | 8 | https://curtain.proteo.info/#/6b02c724-db32-4794-84e3-678c0752e11d | 71 | 40 |
| A549 | 12 | https://curtain.proteo.info/#/68c38f1b-4016-4dd7-b13b-59d925ff09ee | 62 | 26 |
| A549 | 35 | https://curtain.proteo.info/#/9a4fd6bf-a2fd-4579-b6ad-206b2983ca6b | 39 | 14 |
| A549 | 43 | https://curtain.proteo.info/#/47e1361a-6a83-41b4-b8e5-6aa3ca3c1527 | 66 | 21 |
| Brain | 1 | https://curtain.proteo.info/#/3c2e09a1-3e22-4548-9d6f-28379dff6951 | 59 | 19 |
| Brain | 3 | https://curtain.proteo.info/#/330e114b-3f10-44d7-9a68-227bdc5771b9 | 53 | 16 |
| Brain | 7 | https://curtain.proteo.info/#/10d7c884-d02b-4bbf-88e5-08ec74d94e49 | 16 | 4 |
| Brain | 8 | https://curtain.proteo.info/#/1e58aea7-87e1-40f7-8065-7bd87cd6a0c3 | 50 | 9 |
| Brain | 12 | https://curtain.proteo.info/#/d4bc9435-55da-4338-81b2-137aadce7c39 | 15 | 16 |
| Brain | 35 | https://curtain.proteo.info/#/6bf6790b-5985-4eca-a7e0-2e5b9906ae25 | 40 | 91 |
| Brain | 43 | https://curtain.proteo.info/#/d32c7b03-8b4d-403a-8ff5-5028246116f7 | 110 | 54 |

The columns to the right marked -P (non-phosphorylated Rab) and +P (phospho-Rab) list the total number of identified proteins that are enriched in the respective fraction (*p*-value <= 0.05, fold-change FC > 0.6). However, the caveat is that these numbers may include the kinase MST3 (gene STK24) and 'sticky' proteins that are often seen using FLAG-tagged pulldowns of lysates. Therefore, these numbers should be considered as a general guide to the total number of possible specific interactors, as well as non-specific proteins pulled down as a consequence of the specific protocols (e.g., stringency of IP wash) used in this study.
