## Supplementary material for "Defining phospho-Rab signalling networks downstream of LRRK2 kinase": Suppl Table 2

| Gene | Protein | Rab | Function | Annotation | CV | PMID |
|---|---|---|---|---|---|---|
| **A549 - pRab** | | | | | | |
| LGALS3;MAC2 | Galectin-3 | pRab1 | Galactose-binding protein regulates multiple cellular processes including adhesion, proliferation, differentiation, and survival; Reportedly interacts with Rab10, co-localises with Rab7 and LAMP1 | | 25.7 | 34112916, 27693506 |
| * RILPL1;RLP1 | RILP-like protein 1 | pRab1 | Effector of LRRK2-phosphorylated Rab8/Rab10; regulates ciliogenesis and centrosome cohesion | * Significantly enriched in pRab3/8 (brain) and pRab1/3/8/12 (A549) | | 29125462, 41314214 |
| * TBC1D2;PARIS1; | TBC1 domain family member 2A | pRab1 | GAP for Rab7A; effector of Rac1; regulates E-cadherin degradation; LRRK1 is recruited to the autolysosome and regulates autophagy by activating TBC1D2 switching off Rab7 signalling | * Significantly enriched in pRab1/3/8/12/35/43 (A549) | | 29599388 |
| * RILPL1;RLP1 | RILP-like protein 1 | pRab3 | Effector of LRRK2-phosphorylated Rab8/Rab10; regulates ciliogenesis and centrosome cohesion | *Significantly enriched in pRab3/8 (brain) and pRab1/3/8/12 (A549) | 35.5, 36.5, 38.8, 49.6, | 34112916, 27693506 |

| Gene | Protein | Rab | Function | Annotation | CV | PMID |
|---|---|---|---|---|---|---|
| * RILPL2;RLP2 | RILP-like protein 2 | pRab3 | Effector of LRRK2-phosphorylated Rab8/Rab10; regulates ciliogenesis and actin dynamics | *Significantly enriched in pRab3/12 (brain) and pRab3/8/12 (A549) | 49.2 | 34112916, 27693506 |
| * SPAG9;HSS; | JNK-interacting protein 4 | pRab3 | Scaffold protein that organises MAPK signalling; adaptor protein that interacts with dynein-dynactin complex or kinesin for lysosomal positioning; interacts with pRab downstream of LRRK2 | *Significantly enriched in pRab3/8 (brain & A549) | 14.6, 16.6 | 29146937, 40990465 |
| * TBC1D2;PARIS1; | TBC1 domain family member 2A | pRab3 | GAP for Rab7A; effector of Rac1; regulates E-cadherin degradation; LRRK1 is recruited to the autolysosome and regulates autophagy by activating TBC1D2 switching off Rab7 signalling | * Significantly enriched in pRab1/3/8/12/35/43 (A549) | | 29599388 |
| CAD | CAD protein | pRab7 | Multifunctional protein; catalyses initial steps in de novo pyrimidine biosynthesis pathway; STRING suggests interaction with tubulin | | 15.2 | 34288185 |

| Gene | Protein | Rab | Function | Annotation | CV | PMID |
|---|---|---|---|---|---|---|
| HPS3 | Hermansky-Pudlak syndrome 3 protein | pRab7 | HPS3/HPS5/HPS6 form the BLOC-2 complex; BLOC-2 regulates function and trafficking of lysosome-related organelles; interacts with Rab32 and Rab38 | | 20.8 | 30945407 |
| HPS5 | Hermansky-Pudlak syndrome 5 protein | pRab7 | BLOC-2 complex subunit | | 18.2 | 30945407 |
| HPS6 | Hermansky-Pudlak syndrome 6 protein | pRab7 | BLOC-2 complex subunit | | 19.6 | 30945407 |
| * OCRL;OCRL1 | Inositol polyphosphate 5-phosphatase OCRL-1 | pRab7 | Hydrolyses PI(4,5)P2 and PI(3,4,5)P3; regulates membrane trafficking at the Golgi, vesicles, endosomal compartments and plasma membrane; regulates ciliogenesis; established Rab effector | *Significantly enriched in Rab1/8 (brain & A549), pRab7/35/43(A549) | 64.8 | 21971085 |
| * PIP4P1;C14ORF9; TMEM55B | Type 1 phosphatidylinositol 4,5-bisphosphate 4-phosphatase | pRab7 | TMEM55B; lysosomal transmembrane protein that regulates cellular stress and lysosomal repair; forms complex with RILPL1 and similar proteins | * * Significantly enriched in Rab1/12/35/43 (brain), Rab1/12/35 (A549), pRab7/8 (A549), pRab7 (brain) | 55.5 | 41314214, 38168055 |

| Gene | Protein | Rab | Function | Annotation | CV | PMID |
|---|---|---|---|---|---|---|
| WDR91;HSPC049 | WD repeat-containing protein 91 | pRab7 | Negative regulator of PI3-kinase on endosomal membranes to regulate fusion and cargo delivery | | 31.3 | 39426373 |
| MLPH;SLAC2A | Melanophilin | pRab8 | Rab27A effector; links melanosomes to MYO5A actin motor for peripheral melanosome transport | | 19.6, 36.7, 39.1, 40.7 | 30699046 |
| * PIP4P1;C14ORF9; TMEM55B | Type 1 phosphatidylinositol 4,5-bisphosphate 4-phosphatase | pRab8 | TMEM55B; lysosomal transmembrane protein that regulates cellular stress and lysosomal repair; forms complex with RILPL1 and similar proteins | * * Significantly enriched in Rab1/12/35/43 (brain), Rab1/12/35 (A549), pRab7/8 (A549), pRab7 (brain) | 17.1 | 41314214, 38168055 |
| PPM1H;ARHCL1 | Protein phosphatase 1H | pRab8 | Metal dependent phosphatase that counteracts LRRK2 activity | *Significantly enriched in pRab3/8/35 (brain), pRab8 (A549) | 44.8 | 31663853, 34580980 |
| * SPAG9;HSS | JNK-interacting protein 4 | pRab8 | Scaffold protein that organises MAPK signalling; adaptor protein that interacts with dynein-dynactin complex or kinesin for lysosomal positioning; interacts with pRab downstream of LRRK2 | *Significantly enriched in pRab3/8 (brain & A549) | 31.2, 42 | 29146937, 40990465 |

| Gene | Protein | Rab | Function | Annotation | CV | PMID |
|---|---|---|---|---|---|---|
| * TBC1D2;PARIS1 | TBC1 domain family member 2A | pRab8 | GAP for Rab7A; effector of Rac1; regulates E-cadherin degradation; LRRK1 is recruited to the autolysosome and regulates autophagy by activating TBC1D2 switching off Rab7 signalling | * Significantly enriched in pRab1/3/8/12/35/43 (A549) | 47.5 | 29599388 |
| TBC1D8B | TBC1 domain family member 8B | pRab8 | Rab11 GAP involved in vesicular recycling | | 32.3 | 30661770, 31732614 |
| * RILPL1;RLP1 | RILP-like protein 1 | pRab8 | Effector of LRRK2-phosphorylated Rab8/Rab10; regulates ciliogenesis and centrosome cohesion | * Significantly enriched in pRab3/8 (brain) and pRab1/3/8/12 (A549) | 45.3, 46.0, 48.1, 48.1 | 34112916, 27693506 |
| * RILPL2;RLP2 | RILP-like protein 2 | pRab8 | Effector of LRRK2-phosphorylated Rab8/Rab10; regulates ciliogenesis and actin dynamics | *Significantly enriched in pRab3/12 (brain) and pRab3/8/12 (A549) | 48.7 | 34112916, 27693506 |
| MOB4;MOB3 | | pRab8 | Component of STRIPAK complexes that regulate signalling pathways including Hippo, MAPK, and cytoskeletal remodelling | | 16.2 | 35737712 |

| Gene | Protein | Rab | Function | Annotation | CV | PMID |
|---|---|---|---|---|---|---|
| ACAP2;CENTB2; | ACAP2 | pRab12 | ARF6 GAP that regulates ARF6-dependent membrane trafficking and cell migration; Rab35 effector | Significantly enriched in Rab8/35/43 (brain), and Rab8/35, pRab12 (A549) | 73.3 | 30905672 |
| EHBP1L1 | EH domain-binding protein 1-like 1 | pRab12 | Rab effector protein that regulates intracellular trafficking and cytoskeletal organisation | | 53.7 | 36754282 |
| KLHL26 | Kelch-like protein 26 | pRab12 | Member of the KLHL family which may be involved in ubiquitination | | 30.7 | 23676014 |
| *OPTN;FIP2; | Optineurin | pRab12 | Multifunctional adaptor protein; functions in Golgi maintenance, exocytosis and trafficking via myosin VI; Rab effector; involved in innate immune response | * Significantly enriched in Rab1/8 (A549 and brain) and pRab12 (A549) | 69.4 | 23357852 |
| * RILPL1;RLP1 | RILP-like protein 1 | pRab12 | Effector of LRRK2-phosphorylated Rab8/Rab10; regulates ciliogenesis and centrosome cohesion | * Significantly enriched in pRab3/8 (brain) and pRab1/3/8/12 (A549) | 20.9, 23.1, 23.6, 27.2 | 34112916, 27693506 |
| * RILPL2;RLP2 | RILP-like protein 2 | pRab12 | Effector of LRRK2-phosphorylated Rab8/Rab10 | enriched in pRab3/12 (brain) and pRab3/8/12 (A549) | 26.9 | 34112916, 27693506 |

| Gene | Protein | Rab | Function | Annotation | CV | PMID |
|---|---|---|---|---|---|---|
| * TBC1D2;PARIS1; | TBC1 domain family member 2A | pRab12 | GAP for Rab7A; effector of Rac1; regulates E-cadherin degradation; LRRK1 is recruited to the autolysosome and regulates autophagy by activating TBC1D2 switching off Rab7 signalling | * Significantly enriched in pRab1/3/8/12/35/43 (A549) | 33.1 | 29599388 |
| MGST3 | Microsomal glutathione S-transferase 3 | pRab35 | Expression linked to neurodegenerative diseases including PD; regulates alpha-synuclein under oxidative stress | * Significantly enriched in Rab43 (brain) and pRab35 (A549 & brain) | 16.2 | 38971310 |
| * OCRL;OCRL1 | Inositol polyphosphate 5-phosphatase OCRL-1 | pRab35 | Hydrolyses PI(4,5)P2 and PI(3,4,5)P3; regulates membrane trafficking at the Golgi, vesicles, endosomal compartments and plasma membrane; regulates ciliogenesis; established Rab effector | *Significantly enriched in Rab1/8 (brain & A549), pRab7/35/43(A549) | 13.7 | 21971085 |

| Gene | Protein | Rab | Function | Annotation | CV | PMID |
|---|---|---|---|---|---|---|
| * TBC1D2;PARIS1; | TBC1 domain family member 2A | pRab35 | GAP for Rab7A; effector of Rac1; regulates E-cadherin degradation; LRRK1 is recruited to the autolysosome and regulates autophagy by activating TBC1D2 switching off Rab7 signalling | * Significantly enriched in pRab1/3/8/12/35/43 (A549) | 23.3 | 29599388 |
| FAM98A | Protein FAM98A | pRab43 | Forms a complex with PLEKHM1, DEF8, and NDEL1, which interacts with RAB7 | | 8.3 | 27777970 |
| IMPA2 | Inositol monophosphatase 2 | pRab43 | Hydrolyses inositol monophosphates | Significantly enriched in pRab43 (A549 & brain) | 13.6 | 36841024 |
| * TBC1D2;PARIS1 | TBC1 domain family member 2A | pRab43 | GAP for Rab7A; effector of Rac1; regulates E-cadherin degradation; LRRK1 is recruited to autolysosome, regulates autophagy by activating TBC1D2 switching off Rab7 signalling | * Significantly enriched in pRab1/3/8/12/35/43 (A549) | 25.7 | 29599388 |
| * OCRL;OCRL1 | Inositol polyphosphate 5-phosphatase OCRL-1 | pRab43 | Hydrolyses PI(4,5)P2 and PI(3,4,5)P3; regulates membrane trafficking at the Golgi, vesicles, ER and PM | *Significantly enriched in Rab1/8 (brain & A549), pRab7/35/43(A549) | 119.5 | 21971085 |

| Brain - pRabs | | | | | | |
|---|---|---|---|---|---|---|
| **Gene** | **Protein** | **Rab** | **Function** | **Annotation** | **CV** | **PMID** |
| KIF2A | Kinesin-like protein KIF2A | pRab3 | Microtubule-dependent kinesin motor | | 36.8, 38.8, 42, 42.5 | |
| * PPM1H | Protein phosphatase 1H | pRab3 | Metal dependent phosphatase that counteracts LRRK2 activity | *Significantly enriched in pRab3/8/35 (brain), pRab8 (A549) | | 31663853, 34580980 |
| * RILPL1 | RILP-like protein 1 | pRab3 | Effector of LRRK2-phosphorylated Rab8/Rab10; regulates ciliogenesis and centrosome cohesion | * Significantly enriched in pRab3/8 (brain) and pRab1/3/8/12 (A549) | 39.9 | 4112916, 27693506 |
| * RILPL2 | RILP-like protein 2 | pRab3 | Effector of LRRK2-phosphorylated Rab8/Rab10; regulates ciliogenesis and actin dynamics | Significantly enriched in pRab3/12 (brain) and pRab3/8/12 (A549) | 41.2 | 4112916, 27693506 |
| SLMAP | Sarcolemmal membrane-associated protein | pRab3 | Forms extended t (SIKE1:SLMAP) STRIPAK complex; reported functions in membrane fusion and vesicle transport | | 47.8 | 29063833 |
| *SPAG9;HSS; | C-Jun-amino-terminal kinase-interacting protein 4 | pRab3 | Scaffold protein that organises MAPK signalling; interacts with dynein-dynactin/kinesin, lysosomal positioning; pRab interactor | *Significantly enriched in pRab3/8 (brain & A549) | 22.9, 26.5 | 31233748 |

| Gene | Protein | Rab | Function | Annotation | CV | PMID |
|---|---|---|---|---|---|---|
| * PIP4P1;C14ORF9;TMEM55B | Phosphatidylinositol 4,5-bisphosphate 4-phosphatase 1 | pRab7 | TMEM55B; lysosomal transmembrane protein; binds RILPL1 at lysosomal membrane recruited by pRabs | * Significantly enriched in Rab1/12/35/43 (brain), Rab1/12/35 (A549), pRab7/8 (A549), pRab7 (brain) | 66.23 | 41314214, 38168055 |
| * PPM1H | Protein phosphatase 1H | pRab8 | Metal dependent phosphatase that counteracts LRRK2 activity | *Significantly enriched in pRab3/8/35 (brain), pRab8 (A549) | 35.35, 35.99 | 31663853, 34580980 |
| * RILPL1 | RILP-like protein 1 | pRab8 | Effector of LRRK2-phosphorylated Rab8/Rab10; regulates ciliogenesis and centrosome cohesion | * Significantly enriched in pRab3/8 (brain) and pRab1/3/8/12 (A549) | 37.51 | 4112916, 27693506 |
| *SPAG9;HSS; | C-Jun-amino-terminal kinase-interacting protein 4 | pRab8 | Scaffold protein, MAPK signalling; adaptor protein, dynein-dynactin complex or kinesin for lysosomal positioning; interacts with pRab downstream of LRRK2 | *Significantly enriched in pRab3/8 (brain & A549) | 17.43, 26.60 | 29146937, 40990465 |
| MT1 | Metallothionein-1 | pRab12 | Bind heavy metals to regulate cell stress | | 29.40 | 34804020 |
| MT2 | Metallothionein-2 | pRab12 | Bind heavy metals regulate cell stress | | 34.61 | 36889441 |

| Gene | Protein | Rab | Function | Annotation | CV | PMID |
|---|---|---|---|---|---|---|
| MT3 | Metallothionein-3 | pRab12 | Binds heavy metals to regulate cell stress; MT3 important in CNS for protective neurons | | 28.32 | 35503771 |
| * RILPL2 | RILP-like protein 2 | pRab12 | Effector of LRRK2-phosphorylated Rab8/Rab10 | enriched in pRab3/12 (brain) and pRab3/8/12 (A549_ | 51.81 | 4112916, 27693506 |
| * PPM1H | Protein phosphatase 1H | pRab35 | Metal dependent phosphatase that counteracts LRRK2 activity | *Significantly enriched in pRab3/8/35 (brain), pRab8 (A549) | 12.11, 13.63 | 31663853, 34580980 |
| RHOT1 | Mitochondrial Rho GTPase 1 | pRab35 | GTPase domains and calcium-binding EF-hand motifs; Anchors mitochondria to motors, PINK1/Parkin substrate | Significantly enriched in Rab43, pRab35 (brain) | 13.07, 14.17 | 24256248 |
| RHOT2 | Mitochondrial Rho GTPase 2 | pRab35 | Anchors mitochondria to motor complexes | | 14.17, 29.33 | 24256248 |
| RIMS3 | Regulating synaptic membrane exocytosis protein 3 | pRab35 | Regulates synaptic membrane exocytosis | | 20.62 | |
| *TXNDC5 | Thioredoxin domain-containing protein 5 | pRab35 | Protein disulfide isomerase; ER | *Significantly enriched pRab35/43 (brain) | 9.73 | 38629066 |
| *SBF2 | Myotubularin-related protein 13 | pRab43 | Myotubularin pseudo-phosphatase, regulates PI(3,5)P2 on endosomes with MTMR2; Reported GEF for Rab21 & Rab28 | *Significantly enriched Rab35, pRab43 (brain), Rab3/35 (A549) | 16.33 | 25648148 |

| Gene | Protein | Rab | Function | Annotation | CV | PMID |
|---|---|---|---|---|---|---|
| TBC1D5 | TBC1 domain family member 5 | pRab43 | TBC-domain GAP for Rab7; associates with retromer to regulate endosomal Rab7 inactivation; binds to retromer complex | | 53.92 | 29667289 |
| *TXNDC5 | Thioredoxin domain-containing protein 5 | pRab43 | Protein disulfide isomerase; localised to ER | *Significantly enriched pRab35/43 (brain) | 32.47 | 38629066 |
| UNC13A | Protein unc-13 homolog A | pRab43 | Synaptic vesicle priming factor (Munc13-1); ALS/FTD risk gene; | *Significantly enriched Rab3/8/12, pRab43 (brain) | 14.50, 17.1 | 42403099 |
| VAMP2 | Vesicle-associated membrane protein 2 | pRab43 | SNARE protein that regulates membrane fusion and neurotransmitter release at the active zone | | 15.63, 24.68 | 36618823 |
| VPS26A | Vacuolar protein sorting-associated protein 26A | pRab43 | Retromer subunit (Retromer complex consists of VPS35-VPS26A-VPS26B-VPS29); endosomal cargo sorting; Rab7 association | | 45.30 | 26199408 |
| VPS26B | Vacuolar protein sorting-associated protein 26B | pRab43 | Retromer subunit; | | 50.38, 51.23 | 26199408 |
| VPS29 | Vacuolar protein sorting-associated protein 29 | pRab43 | Retromer subunit | | 46.16 | 26199408 |
| VPS35 | Vacuolar protein sorting-associated protein 35 | pRab43 | Retromer subunit; VPS35-D620N causes PD (late onset) | | 50.00 | 39197569 |

### Supplementary Table 2

List of potential interactors that were enriched with phosphorylated Rab GTPases from A549 and mouse brain lysates. The PMID column are citations that correspond to PubMed identification numbers. The CV column denotes the Coefficient of Variation for each replicate group. **GREEN** are values below 20 indicating strong correlations within the group. **ORANGE** are values between 20-50, while **RED** indicate values above 50, which are the weakest correlations.
